# Functional Localization of the Human Auditory and Visual Thalamus Using a Thalamic Localizer Functional Magnetic Resonance Imaging Task

**DOI:** 10.1101/2024.04.28.591516

**Authors:** John C. Williams, Philip N. Tubiolo, Zu Jie Zheng, Eilon B. Silver-Frankel, Dathy T. Pham, Natalka K. Haubold, Sameera K. Abeykoon, Anissa Abi-Dargham, Guillermo Horga, Jared X. Van Snellenberg

**Affiliations:** Department of Psychiatry and Behavioral Health, Renaissance School of Medicine at Stony Brook University, Stony Brook, NY 11794; Department of Biomedical Engineering, Stony Brook University, Stony Brook, NY 11794; State University of New York Downstate Health Sciences University College of Medicine, Brooklyn, NY 11203; Department of Psychiatry, Columbia University Vagelos College of Physicians and Surgeons, New York-Presbyterian / Columbia University Irving Medical Center, New York, NY 10032; New York State Psychiatric Institute, New York, NY 10032; Department of Neurobiology and Behavior, Cornell University, Ithaca, NY 14853; Department of Radiology, Renaissance School of Medicine at Stony Brook University, Stony Brook, NY 11794; Department of Psychology, Stony Brook University, Stony Brook, NY 11794

**Keywords:** Auditory processing, visual processing, medial geniculate, lateral geniculate, functional localizer task, resting-state functional connectivity

## Abstract

Functional magnetic resonance imaging (fMRI) of the auditory and visual sensory systems of the human brain is an active area of investigation in the study of human health and disease. The medial geniculate nucleus (MGN) and lateral geniculate nucleus (LGN) are key thalamic nuclei involved in the processing and relay of auditory and visual information, respectively, and are the subject of blood-oxygen-level-dependent (BOLD) fMRI studies of neural activation and functional connectivity in human participants. However, localization of BOLD fMRI signal originating from neural activity in MGN and LGN remains a technical challenge, due in part to the poor definition of boundaries of these thalamic nuclei in standard T1-weighted and T2-weighted magnetic resonance imaging sequences. Here, we report the development and evaluation of an auditory and visual sensory thalamic localizer (TL) fMRI task that produces participant-specific functionally-defined regions of interest (fROIs) of both MGN and LGN, using 3 Tesla multiband fMRI and a clustered-sparse temporal acquisition sequence, in less than 16 minutes of scan time. We demonstrate the use of MGN and LGN fROIs obtained from the TL fMRI task in standard resting-state functional connectivity (RSFC) fMRI analyses in the same participants. In RSFC analyses, we validated the specificity of MGN and LGN fROIs for signals obtained from primary auditory and visual cortex, respectively, and benchmark their performance against alternative atlas- and segmentation-based localization methods. The TL fMRI task and analysis code (written in Presentation and MATLAB, respectively) have been made freely available to the wider research community.

## 1 Introduction

The thalamus is a key neural processing hub in human sensory perception (Kandel, 2013; Mai & Paxinos, 2012; Sherman, 2016; Ward, 2013). Disruption of thalamocortical connectivity has been implicated in the pathogenesis of multiple neurological and psychiatric diseases (Anticevic et al., 2014; Anticevic et al., 2015; Chun et al., 2017; Chun et al., 2014; Giraldo-Chica et al., 2018; Karlsgodt, 2016; Kelly et al., 2018; Kim et al., 2014; Kubota et al., 2013; Murray & Anticevic, 2017; Pasternak et al., 2018; Peters et al., 2016; Pettersson-Yeo et al., 2011; Sheffield et al., 2020; Tamnes & Agartz, 2016; Tu et al., 2019; Woodward & Cascio, 2015; Woodward et al., 2017; Woodward & Heckers, 2016; Woodward et al., 2012), including potential specific involvement of sensory thalamic nuclei (Chun et al., 2017; Chun et al., 2014; Giraldo-Chica et al., 2015; Huang et al., 2024). Accordingly, the study of neural activity in sensory thalamic nuclei and their functional connectivity with sensory cortex is an area of active investigation in the study of the human brain in health and disease (Almasabi et al., 2022; Bartlett, 2013; Chun et al., 2017; Chun et al., 2014; Erskine et al., 2016; Papadopoulou et al., 2019; Schmid et al., 2010; Truong et al., 2015), including through the use of blood-oxygen-level-dependent (BOLD) functional magnetic resonance imaging (fMRI) of human participants.

Within the thalamus, the medial geniculate nucleus (MGN) and lateral geniculate nucleus (LGN) are integral components of the human auditory and visual pathways, respectively, and play critical roles in the modulation and relay of incoming sensory information to the cortex (Kandel, 2013; Sherman, 2016; Ward, 2013). The MGN receives incoming sensory information from the cochlear nucleus, primarily through the inferior colliculus and superior olivary complex, and provides ascending inputs to the auditory cortex (AC; Bartlett, 2013; Brodal, 1981; Kandel, 2013; Lee, 2013). The AC, in turn, projects distally back to MGN, inferior colliculus, and superior olive (Bartlett, 2013; Lee, 2013), in order to facilitate integration of modulatory and contextual information, such as prediction error signals (Li & Ebner, 2016; Malmierca et al., 2015; Parras et al., 2017). Similarly, the LGN receives inputs from the retina, as well as the superior colliculus, and provides inputs to the visual cortex (VC), which projects back to LGN and superior colliculus (Gilbert & Li, 2013; Lee et al., 2010; Sherman, 2016).

BOLD fMRI studies that aim to measure activity within MGN and LGN hinge upon accurate localization of these structures. However, identifying regions of interest (ROIs) within the thalamus that contain BOLD activation patterns specific to auditory and visual perception using standard atlases or segmentation techniques is hindered by the relatively small size of these nuclei (e.g., estimates of approximately 120 mm^3^ and 60 mm^3^, respectively, for LGN (Muller-Axt et al., 2021) and MGN (Garcia-Gomar et al., 2019)), and a lack of distinct anatomical landmarks in the posterior thalamus that can be identified using T1-weighted (T1w) and T2-weighted (T2w) imaging sequences that causes segmentation algorithms to rely heavily on priors (relative to individual-specific anatomical information). Moreover, individual variability in the location and size of nuclei (Andrews et al., 1997; Garcia-Gomar et al., 2019; Giraldo-Chica & Schneider, 2018; Kiwitz et al., 2022; Rademacher et al., 2002) is exacerbated by both the difficulty of precise spatial normalization to standardized spaces, and the resulting imprecision in the coregistration between anatomical structural images and echo planar images (EPIs) measuring hemodynamic BOLD signal.

One broadly successful approach to localizing brain regions with specific neural functions that are otherwise difficult to localize based on anatomical scans alone is to develop and utilize a task that specifically elicits the function subserved by the targeted neural region, commonly known as a functional localizer task (Berman et al., 2010; Dodell-Feder et al., 2011; Fedorenko et al., 2010; Jacoby et al., 2016; Jiang et al., 2013; Kastner et al., 2004; Saxe et al., 2006). Consequently, BOLD fMRI sensory thalamic localizer (TL) tasks have been developed to measure human sensory thalamic responses *in vivo*, enabling group-wide and/or single-participant localization of geniculi experimentally (Chen et al., 1998; Denison et al., 2014; Hess et al., 2009; Jiang et al., 2013; Kastner et al., 2004; Schneider & Kastner, 2009; Sitek et al., 2019).

In this work, we developed and implemented an auditory and visual sensory TL fMRI task, using high-resolution multiband fMRI (Moeller et al., 2010), with the aim of producing functionally-defined ROIs (fROIs) optimal for extracting information from other BOLD images acquired from the same participant. To improve functional specificity, this task uses clustered-sparse temporal acquisition (Schmidt et al., 2008; Yang et al., 2000; Zaehle et al., 2007), through which volumes are collected only immediately after the presentation of task stimuli, thereby reducing the confounding effect of concurrent scanner noise, which is critical when studying auditory processing with fMRI (Bandettini et al., 1998; Schmidt et al., 2008). The task developed here builds on prior sensory TL tasks by 1) combining both visual and auditory stimulation in a single task to simultaneously identify fROIs for MGN and LGN; 2) leveraging multiband acceleration to improve spatial resolution and increase the number of data points (volumes) acquired with each acquisition cluster, which in turn facilitates 3) using a shorter (∼15 minute) total task length while still obtaining high quality fROIs; and 4) is developed and tested at 3 Tesla (as opposed to higher fields) in order to maximize availability to investigators.

We report the utility of this sensory TL task in producing participant-level MGN and LGN fROIs, validate their specificity for auditory and visual processing using resting-state (RS) fMRI scans acquired from the same participants, and demonstrate the performance of these fROIs relative to widely used atlas and segmentation-based techniques. Code for the TL fMRI task employed here, as well as participant-level analysis code for generating MGN and LGN fROIs, are also made publicly available for use by the wider research community.

## 2 Materials and Methods

### 2.1 Overview

This is a two-site study of healthy participants who completed task procedures at either the New York State Psychiatric Institute (NYSPI) or Stony Brook University (SBU). All research procedures at NYSPI were approved by the New York State Psychiatric Institute Institutional Review Board; all research procedures at SBU were approved by the Stony Brook University Institutional Review Board. Participants were recruited with advertisements and all individuals provided written consent prior to their participation.

### 2.2 General Inclusion and Exclusion Criteria

Data at each site were collected from healthy adult participants, ages 18-55, over the course of other studies, and thus inclusion and exclusion criteria varied slightly between sites. Full inclusion and exclusion criteria for each site are described in the ***Inclusion and Exclusion Criteria*** section of **Supplementary Materials and Methods.** All participants were free of any major neurological disorders, psychiatric disorders, current substance use disorders, and hearing impairment. Participants at SBU were additionally tested for normal hearing thresholds and speech recognition for monosyllabic words. Psychiatric diagnoses were determined at NYSPI according to the Diagnostic and Statistical Manual of Mental Disorders, Fourth Edition, Text Revision (DSM-IV-TR; American Psychiatric Association, 2000), and at SBU according to the Diagnostic and Statistical Manual of Mental Disorders, Fifth Edition (DSM-5; American Psychiatric Association, 2013).

### 2.3 Demographic and Clinical Assessments

Handedness was assessed using the Edinburgh Handedness Inventory (EHI; Oldfield, 1971), and socioeconomic status using the Hollingshead Four Factor Index of Socioeconomic Status (Hollingshead, 1975). Family history of mental disorders was assessed using the Family History Screen (FHS; Milne et al., 2009). Clinical measures were assessed at NYSPI using the Structured Clinical Interview for the DSM-IV Axis I Disorders (SCID-I; First & Gibbon, 2004), and at SBU using the Structured Clinical Interview for DSM-5, Research Version (SCID-5-RV; First et al., 2015), and at both sites using the Positive and Negative Syndrome Scale (PANSS; Kay et al., 1987). At SBU, hearing thresholds were evaluated either using a GSI Automated Method for Testing Auditory Sensitivity Pro (Grason-Stadler, Eden Prairie, Minnesota) automated audiometry system, or by a licensed audiologist or otolaryngologist, and speech recognition for monosyllabic words using the Northwestern University Auditory Test No. 6 (Carhart & Tillman, 1966).

### 2.4 fMRI Acquisition

Prior to magnetic resonance (MR) scanning procedures, participants completed an MRI clearance form to assess for metallic implants and past experiences with metal, and were administered a urine toxicology test, as well as a urine pregnancy screen for biologically female participants. MR scanning was carried out using a 3 Tesla General Electric (GE; Boston, MA) MR750 with a NOVA 32-channel head coil at NYSPI, and a 3 Tesla Siemens (Munich, Germany) MAGNETOM Prisma at the SBU Social, Cognitive, and Affective Neuroscience (SCAN) Center with a Siemens 64-channel head-and-neck coil (Keil et al., 2013). All T2* BOLD EPI data were acquired in alternating, opposing phase-encode (PE) directions, i.e., two anterior-posterior (AP) PE runs and two posterior-anterior (PA) PE runs.

The following data were acquired during each session at SBU: 1) high-resolution (0.80 mm isotropic voxels) anatomical T1w and T2w scans; 2) four T2*-weighted BOLD EPI RS runs; 3) four T2*-weighted BOLD EPI TL runs; 4) two spin echo field maps, one AP and one PA; and 5) one B0 field map. BOLD fMRI data were acquired at NYSPI with a TR of 850 ms and field of view of 192 mm, while BOLD data acquired at SBU were acquired with a TR of 800 ms and field of view of 204 mm. BOLD data were acquired from both sites with 2mm isotropic voxels, a flip angle of 60 degrees, echo time of 25 ms, and multiband factor of 6. The duration of each RS run was 7 minutes 34 seconds at NYSPI and 7 minutes 38 seconds at SBU. At both sites (NYSPI and SBU), each thalamic localizer run was 3 minutes 46 seconds in duration. Participants were asked to complete 4 runs each of the TL fMRI task and RS fMRI, with a minimum of two runs of each required for study completion and inclusion in the final analyzed dataset.

### 2.5 Sensory Thalamic Localizer fMRI Task Procedures

The goal of the sensory thalamic localizer task is to identify regions of the thalamus specifically responsive to auditory and visual stimuli in each participant, as has been done in prior work (Jiang et al., 2013; Kastner et al., 2004). This task uses clustered-sparse temporal acquisition (Schmidt et al., 2008; Yang et al., 2000; Zaehle et al., 2007), in which fMRI volumes are not acquired while task stimuli are presented. Instead, fMRI is acquired immediately after the presentation of stimuli, in order to minimize the effect of scanner noise from BOLD data acquired; otherwise, the sound produced by the MR scanner while volumes are being acquired could confound detected responses to stimuli (Schmidt et al., 2008). In this task, 3 volumes of BOLD fMRI are acquired at the end of each trial. Acquisition clusters were timed such that a) the hemodynamic response to scanner noise produced by the prior acquisition cluster should have returned to baseline levels, and b) the hemodynamic response to stimuli presented between two clusters is at or near its peak when the acquisition cluster is acquired.

A schematic diagram of the sensory TL fMRI task is shown in **Figure 1**. In each run **(Figure 1A)**, participants are presented with 16 trials of either visual or auditory stimulation in a pseudorandom order. Each trial consists first of a “gap period” of silence, 9 seconds of stimulation (sufficient to reach the peak of the hemodynamic response in sensory cortex; Inan et al., 2004; Jiang et al., 2013), a second “gap period” of silence equal in duration to the first, and then the clustered acquisition of 3 volumes of BOLD data (TR × 3 in duration). The duration of gap periods is calibrated such that the total trial time is 12 seconds in both NYSPI and SBU data, where the acquisition clusters last 3 TRs. In NYSPI data, this gap period is 225 ms (acquisition cluster duration of 2.55 s); in SBU data, this gap period is 300 ms (acquisition cluster duration of 2.4 s).

**Figure 1.**
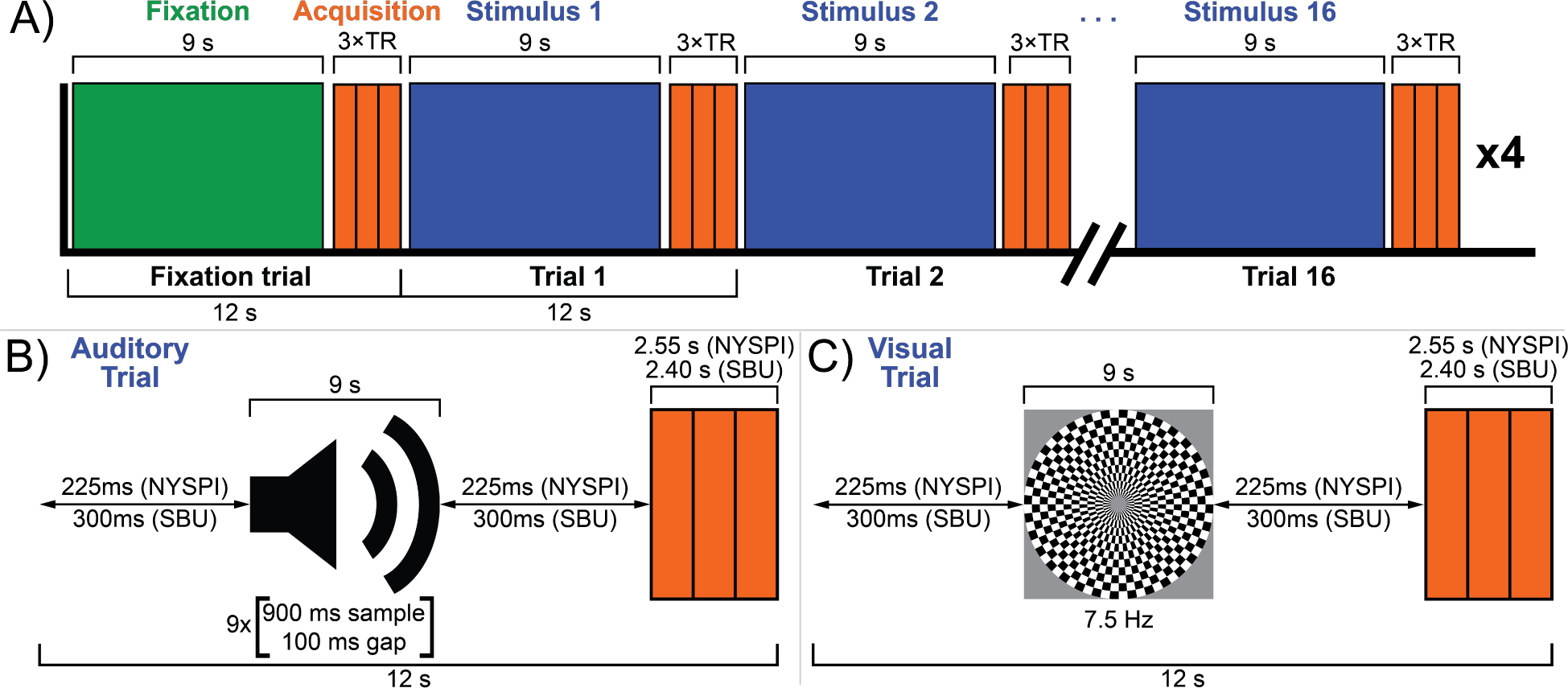
The sensory thalamic localizer (TL) task. A schematic diagram of a TL task run is shown in panel. **A**. Each run begins with a single 12-second fixation trial (green), followed by sixteen 12-second stimulus trials, consisting of 8 auditory and 8 visual trials (blue), presented in pseudorandom order, using clustered-sparse temporal acquisition. Sample auditory and visual trials are respectively shown in panels **B** and **C.** Each trial consists of 9 seconds of stimulation, followed by a single acquisition cluster of 3 volumes (orange). Acquisition clusters are 3 TRs in duration, totaling 2.55 s in NYSPI data and 2.4 s in SBU data. Stimuli are immediately preceded and succeeded by gaps of equal duration, during which no stimulus is presented, in order to bring the total trial time to 12 seconds; this gap time is 225 ms in NYSPI data and 300 ms in SBU data. Auditory trials consist of nine sets of 900 ms amplitude-normalized snippets of instrumental music presented in pseudorandom order and separated by 100 ms gaps. Visual trials consist of a circular checkerboard alternating between black and white at 7.5 Hz. Participants were asked to complete a total of 4 task runs.

Auditory stimulation was presented as nine sets of 900 ms segments of music, followed by 100 ms of silence, in pseudorandom order **(Figure 1B),** similar to prior work (Jiang et al., 2013), in order to reduce neural responses specific to the experience of listening to music, delivered binaurally using a SereneSound headset and transducer (Resonance Technology, Inc., Northridge, CA). Musical segments were obtained from the instrumental song, *Transmission94 (Parts 1 & 2)* by *Bonobo.* This song was chosen to meet the following criteria: 1) instrumental music, to avoid evoking responses specific to speech processing in segments containing perceptible speech fragments; 2) no breaks or pauses in the music; and 3) a long enough song to produce at least 7.5 minutes of audio segments after discarding the first and last 4.8 seconds of the song (to avoid fade in/fade out effects or segments in which the music begins late or ends early; also note that less than 7.5 minutes was ultimately used in the task version reported here). Musical segments were normalized to have the same mean amplitudes in order to avoid sudden changes in amplitude when the presentation switches from one 900 ms segment to another (which may have been selected from disparate portions of the song).

Visual stimulation was presented via a projector with a 60 Hz refresh rate as a circular black-and-white checkerboard with a central fixation cross, alternating between light and dark with a reversal frequency of 7.5 Hz **(Figure 1C),** which was found in prior work to reliably activate both LGN and VC (Kastner et al., 2004; Kastner et al., 2006; Rosa et al., 2010). The presentation of task stimuli was controlled using Presentation software (Version 22.1, Build 10.23.20, Neurobehavioral Systems, Inc., Berkeley, CA, www.neurobs.com) installed on a dedicated laptop. The sensory TL task Presentation code (Williams et al., 2024) is freely available from the Neurobehavioral Systems Archives of Neurobehavioral Experiments and Stimuli at http://www.neurobs.com/ex_files/expt_view?id=302.

### 2.6 Resting-State fMRI Procedures

During RS scans, participants were instructed to look ahead at a fixation cross, remaining still and awake with their eyes open. One of the participants’ eyes was monitored by a research coordinator via an eye tracking camera (although eye tracking was not collected), and eye closure times were recorded to exclude times from analysis times during which participants may have been sleeping, as participants entering into and out of sleep states has been shown to measurably impact on measured RSFC (Tagliazucchi & Laufs, 2014).

### 2.7 fMRI Pre-Processing and Post-Processing

All fMRI data were preprocessed through the following steps of the Human Connectome Project Minimal Preprocessing Pipelines (HCP MPP; Glasser et al., 2013), version 4.2.0 (https://github.com/Washington-University/HCPpipelines): 1) *PreFreeSurfer*, 2) *FreeSurfer*, 3) *PostFreeSurfer,* and 4) *fMRIVolume*. Surface registration was completed by cortical surface matching (MSMsulc; Robinson et al., 2018; Robinson et al., 2014). Preprocessed structural (T1w) and functional (BOLD) images were obtained from *fMRIVolume* normalized to Montreal Neurological Institute (MNI) 152 non-linear 6th-generation space (MNI152NLin6; Grabner et al., 2006).

Nuisance signals for white matter (WM) and cerebro-spinal fluid (CSF) were calculated in unsmoothed data as the average signal in all compartment voxels that remain after an iterative erosion procedure (Power et al., 2014). In this iterative erosion procedure, masks are eroded up to three times, as long as one additional erosion would result in at least two remaining voxels. Six vectors of motion parameters (MPs), the estimated framewise motion in each of the 3 directions of translation and 3 directions of rotation, was also obtained from the HCP MPP. TL and RS BOLD fMRI data were smoothed using a 4 mm full-width-half-maximum Gaussian filter (note that unsmoothed data was retained and used in analyses where noted below).

#### 2.7.1 Resting-State fMRI Data Post-Processing

RS images (both smoothed and unsmoothed) were mode 1000 normalized (divided by the modal value of all in-brain voxels and multiplied by 1000; Power et al., 2012), linearly detrended, and mean centered. Volumes acquired during periods of excess participant motion were removed using volume censoring. Volume censoring (Power et al., 2012; Power et al., 2014; Power et al., 2015; Smyser et al., 2011; Williams et al., 2022) was performed using study-wide thresholds for participant motion, measured by low-pass-filtered framewise displacement (LPF-FD), and run-wise thresholds for whole-brain signal fluctuation, using generalized extreme value low-pass filtered temporal derivative root-mean-squared over voxels (GEV-DV) thresholding, as has been described in detail elsewhere (Williams et al., 2022). Censoring thresholds were determined using resting-state data available for participants with usable TL MGN and LGN fROIs using the Multiband Censoring Optimization Tool (Williams et al., 2022) (https://github.com/CNaP-Lab/MCOT/), resulting in an LPF-FD threshold (*Φ_F_*) of 0.07587 mm and GEV-DV parameter (*d_G_*) of 3.105 (arbitrary units).

Volumes acquired at times during which a participant eye closure lasted longer than 3 seconds in duration were additionally removed through volume censoring, as were volumes acquired between eye closure periods that occurred less than 30 seconds apart. Contiguous clusters of data that were less than 8 s in duration after censoring were discarded, as were resting state runs with fewer than 1.5 minutes of remaining data. Participants with fewer than either 2 runs of data remaining, or fewer than 5 minutes of remaining data in total, were excluded from resting-state analyses.

RS runs were filtered using a 0.009-0.08 Hz band-pass second-order zero-phase Butterworth filter, with censored time points replaced by linear interpolation prior to band-pass filtering before being discarded from analysis. The first and last 22 seconds of each time series were discarded to remove discontinuity (edge) artifacts present after band-pass filtering; approximately 22 seconds of data were removed from the start and end of each run (slightly different between sites due to the different TRs: 22.1 s / 26 volumes at NYSPI, 22.4 s / 28 volumes at SBU).

### 2.8 Sensory Thalamic Localizer fMRI Task Analysis

#### 2.8.1 Overview

The objective in analyzing the TL task is to identify voxels within defined regions of posterior thalamus that show coactivation with either auditory cortex or visual cortex, but not both, over the course of alternating auditory and visual stimulation. To achieve this, TL task data were analyzed to first identify a cluster within each hemisphere’s auditory cortex (AC) and visual cortex (VC) that was maximally responsive to either auditory or visual stimulation, respectively (section **2.8.3**). Average AC and VC time series were extracted from these clusters (section **2.8.4**).

Participant-specific hemispheric thalamic search region (TSR) masks for MGN and LGN (MGN-TSR and LGN-TSR, respectively) were generated (section **2.8.2**). Voxels within the MGN-TSR and LGN-TSR were assessed for coactivation with the AC and VC time series during the TL task (using Pearson’s partial correlations; see section **2.8.5)** and thresholded based on this coactivation. Voxels showing coactivation with both AC and VC time series were removed to improve specificity (section **2.8.5)**. The largest remaining contiguous cluster within each hemisphere’s MGN-TSR and LGN-TSR were extracted as each participant’s MGN and LGN, respectively.

#### 2.8.2 *A Priori* Regions of Interest

Auditory cortex (AC) and visual cortex (VC) search region masks were generated from Brodmann areas (BAs) using the Wake Forest University PickAtlas Toolbox (WFU; Maldjian et al., 2004; Maldjian et al., 2003). A temporal lobe AC mask was produced from BAs 41 and 42, and an occipital lobe VC mask from BAs 17 and 18, and each was finally dilated in 3D by one voxel.

Hemispheric search region masks for auditory thalamus (MGN-TSR) and visual thalamus (LGN-TSR) were generated for each participant by first applying the FreeSurfer thalamic nuclei segmentation (Iglesias et al., 2018) to each participant’s anterior commissure-posterior commissure (AC-PC) aligned, readout distortion and bias field corrected T1w image. The resulting segmentation output image was warped to MNI152NLin6 space using the warps calculated by the HCP MPP (*acpc_dc2standard.nii.gz*) and resampling to BOLD space, using Connectome Workbench 1.5.0 (Marcus et al., 2011) (*wb_command -volume-resample*). To create an appropriately large search space in posterior thalamus, FreeSurfer estimates of MGN were 3D dilated 3 voxels for the MGN-TSR, and FreeSurfer estimates of LGN were 3D dilated by 1 voxel for the LGN-TSR (as the FreeSurfer LGN ROIs were slightly larger).

To help ensure that fROIs were distinct from surrounding structures, TSR voxels estimated to comprise part of surrounding structures were removed to improve specificity and robustness across participants as follows. First, masks of pulvinar and mediodorsal nuclei were obtained from the FreeSurfer thalamic segmentation. Next, the following masks were obtained from the FreeSurfer segmentation atlas (Desikan et al., 2006; Fischl, 2012; Fischl et al., 2004) output by the HCP MPP (*Atlas_wmparc.2.nii.gz*): cortical gray matter, parahippocampal gyrus WM, insular cortex WM, choroid plexus, pulvinar, mediodorsal nucleus, putamen, pallidum, and hippocampus. The masks of mediodorsal nucleus, insular cortex WM, choroid plexus, pulvinar, mediodorsal nucleus, putamen, and pallidum were 3D dilated by 1 voxel. All voxels overlapping with masks of surrounding structures were then removed from MGN-TSR and LGN-TSR masks. Finally, the posterior-most 2 slices of each TSR mask were then removed to remove voxels in close proximity to midbrain, and any MGN-TSR voxels superior to the inferior-most slice of the pulvinar in either hemisphere were removed to constrain the search region to the inferior aspect of posterior thalamus and remove remaining voxels posterior to the pulvinar FreeSurfer ROI. Probability maps of final MGN-TSR and LGN-TSR masks are shown in **Figure S1.**

#### 2.8.3 Generalized Linear Modeling and Generation of Participant-Level Contrasts

Smoothed TL task fMRI data were analyzed using a generalized linear modeling framework in SPM12 (Friston et al., 2007; Friston, Holmes, et al., 1994; Friston, Jezzard, et al., 1994), version 7771, in MATLAB R2018a (The MathWorks, Inc., Natick, MA), with betas estimated from a design matrix that contained, for each run, a column each for intercept, auditory stimulation, visual stimulation, 6 MPs and their squares, WM signal, and CSF signal, as well as two additional regressors to model the second and third volumes of each cluster. These volume-specific regressors were employed to regress out the effect of T1-relaxation on BOLD signal, as the clustered-sparse temporal acquisition scheme meant that BOLD data were not at steady-state during acquisition (Horga et al., 2014; Schmidt et al., 2008; Zaehle et al., 2007). Task regressors indicating trial type (auditory or visual) were not convolved with a hemodynamic response function (HRF), because the sparse temporal sampling precludes a true timeseries HRF model, as acquisition of volumes is not continuous.

Contrast images of activation from auditory stimulation versus visual stimulation (Auditory − Visual) were then generated for each participant. Each hemisphere’s AC search region masks were thresholded to include only voxels with the top 10% of [Auditory − Visual] contrast values, and VC search region masks were similarly thresholded to include only the bottom 10% of [Auditory − Visual] contrast values. Clusters smaller than 10 contiguous voxels in size were then removed. The single, contiguous cluster within each AC mask and VC mask with the greatest magnitude peak contrast value was used for extraction of AC and VC task time series.

#### 2.8.4 Extraction of Time Series

To obtain AC and VC time series for each participant, TL task BOLD activation was averaged within each hemisphere’s AC and VC clusters, as identified in section **2.8.3**. These hemispheric AC and VC time series were then averaged across hemispheres, resulting in single AC and VC time series for each participant.

Local WM signal was additionally obtained for each TSR for use as nuisance regressors by first identifying WM in the FreeSurfer segmentation atlas located between 1 and 5 voxels from the TSR in any direction, and then extracting average time series from these voxels from unsmoothed BOLD data. Brain-wide gray matter signal was extracted by eroding the FreeSurfer segmentation atlas by one voxel and spatially averaging signal from unsmoothed BOLD data within the resultant mask.

#### 2.8.5 Task Coactivation Analysis and Identification of MGN and LGN Regions of Interest

We aimed to identify thalamic voxels in each TSR that coactivate, over the course of auditory and visual stimulation, with AC and VC. Note that this is distinct from task connectivity analysis methods such as general functional connectivity (Elliott et al., 2019) in that, rather than exploring connectivity after removing task effects, we aimed instead to identify shared patterns of task responsiveness (coactivation). This was done to capitalize on shared variance (connectivity) between sensory cortex and thalamic nuclei across volumes within a task condition, rather than merely identifying thalamic voxels that activate to task. That is, using only a task contrast [Auditory – Visual] would only pick up on differences driven by the task condition, while the coactivation method employed can be driven by these differences in addition to functional connectivity to sensory cortex that is detectable within each fMRI volume acquired within each condition.

Within each TSR (left and right MGN-TSR and LGN-TSR), coactivation between each voxel’s time series and activation in AC and VC (see section **2.8.4**) was estimated using partial correlations, resulting in AC and VC coactivation maps. Note that this produces separate AC and VC coactivation maps for each TSR-MGN and each TSR-LGN search region. Partial correlations included nuisance regressors for average WM, CSF, gray matter signal, local WM (see section **2.8.4**), and a regressor for the second and third image of each cluster.

AC and VC coactivation maps were then thresholded within each TSR to select the highest-ranking voxels (i.e., voxels with the greatest Pearson correlations). TSR-MGN and TSR-LGN upper percentile thresholds were determined separately for each participant, based on the average number of voxels comprising each TSR-MGN and TSR-LGN across hemispheres. The upper percentile threshold for TSR-MGN coactivation maps was 32 divided by the average number of voxels, and the upper percentile threshold for TSR-LGN coactivation maps was 20 divided by the average number of TSR-LGN voxels. These parameter values are arbitrary, and were selected in order to produce reasonably sized and shaped fROIs, with a stricter threshold in the larger TSR-LGN to accommodate overall greater LGN-VC observed coactivation values. Any voxels remaining in both AC and VC coactivation maps of any TSR after thresholding were then removed from both, in order to improve the specificity of resulting fROIs for sensory modality. The largest contiguous cluster remaining within each hemi-thalamus’s TSR-MGN-AC coactivation map, and TSR-LGN-VC coactivation map, were then identified as MGN and LGN, respectively.

Each MGN and LGN fROI was evaluated by multiple members of the study team (J.C.W., P.N.T., and Z.J.Z., and J.X.V.S.) by overlaying them over each participant’s high-resolution T1w anatomical images and temporally averaged BOLD TL task EPIs. If issues were noted by any reviewer, they were examined by two members of the study team (J.C.W. and J.X.V.S.) and a consensus was reached. We excluded fROIs that were considered to be anatomically implausible (e.g., an LGN that is either inferior or medial to MGN, or excessive asymmetry between left and right hemispheric MGN or LGN fROIs); data collected from participants with any fROI that failed this quality check (QC) procedure were excluded from further analyses.

### 2.9 Resting-State Data Analysis

#### 2.9.1 Overview

After localization using the sensory TL task, functional connectivity from MGN and LGN fROIs was evaluated in resting state data obtained from the same participants. Specificity for connectivity with auditory and visual regions was determined by comparing connectivity between primary AC and VC ROIs (obtained independently of the TL task), and compared with connectivity from alternative MGN and LGN ROIs derived from a standardized atlas, the WFU PickAtlas Toolbox (Maldjian et al., 2004; Maldjian et al., 2003), and two thalamic segmentations using T1w images, the FreeSurfer thalamic nuclei segmentation (Iglesias et al., 2018; Su et al., 2019) and the Thalamus Optimized Multi Atlas Segmentation (THOMAS) segmentation (Iglesias et al., 2018; Su et al., 2019) (see section **2.9.2**, below). We then performed whole-brain seed connectivity analyses to determine functional relationships between signals within TL MGN and LGN fROIs, as well as the differences between functional connectivity maps derived from TL ROIs and those alternative ROIs obtained from the aforementioned atlas and T1w segmentations (Iglesias et al., 2018; Maldjian et al., 2004; Maldjian et al., 2003; Su et al., 2019).

#### 2.9.2 *A Priori* Regions of Interest

ROIs for primary AC and primary VC were obtained from the FreeSurfer Desikan-Killiany segmentation atlas (Desikan et al., 2006; Fischl, 2012; Fischl et al., 2004) output by the HCP MPP (*Atlas_wmparc.2.nii*.gz). AC was created from the transverse temporal gyrus parcel (#1034 and #2034), dilated by one voxel in 3D. The VC mask was created from the pericalcarine gray matter parcels (#1021 and #2021), 3D dilating once, removing any voxels not considered to be in gray matter, and then 3D dilating two more times. These ROIs are shown in **Figure S2.** As a benchmark for the potential for bias in RSFC analysis that would be produced by excessive overlap between primary AC and primary VC ROIs described here and the fROIs derived from the TL task, we additionally calculated the Dice-Sørensen similarity coefficient (Dice, 1945; Serrano-Sosa et al., 2021) for each participant’s TL fROI and *a priori* anatomical ROI, bilaterally, in AC and VC separately.

TL MGN and LGN fROIs were obtained from the thalamic localizer task as described above (see section **2.8.5**). An alternative atlas-based set of left and right MGN and LGN ROIs were obtained from the WFU PickAtlas Toolbox (Maldjian et al., 2004; Maldjian et al., 2003); alternative T1w-segmentation based MGN and LGN ROIs for comparison to the thalamic localizer ROIs developed here were obtained from the FreeSurfer thalamic nuclei segmentation (Iglesias et al., 2018) and the Thalamus Optimized Multi Atlas Segmentation (THOMAS) segmentation (Su et al., 2019), shown in **Figure S3.**

#### 2.9.3 Resting-State Functional Connectivity Analysis

For each run, time series were extracted for each ROI (including fROIs) by spatially averaging unsmoothed volumetric resting-state data within ROI voxels. MGN-AC connectivity was estimated by calculating pairwise Pearson’s partial correlations between left MGN and left AC, left MGN and right AC, right MGN and left AC, and right MGN and right AC; these four values were then averaged within each run to create a single run-level MGN-AC connectivity estimate. Both ipsilateral and contralateral connectivity was evaluated in order to account for bilateral processing of sensory stimuli and commissural interhemispheric communication between sensory cortices (Bamiou et al., 2007; Chen et al., 2018; Engel et al., 1991; Ito et al., 2016; Jones, 2007; Lee, 2013; Lee & Winer, 2008; Ramachandra et al., 2020; van der Knaap & van der Ham, 2011), as well as to maximize power. This process was then repeated to additionally calculate MGN-VC, LGN-AC, and LGN-VC connectivity for each run. This procedure was performed for MGN and LGN fROIs obtained from the sensory TL task, as well as the three alternative ROIs (WFU PickAtlas, FreeSurfer, and THOMAS). Differences between MGN-AC and LGN-VC connectivity estimates produced using TL fROIs and those produced using these alternative ROIs were then estimated for each participant.

Volumetric whole-brain seed connectivity images for each hemisphere’s MGN and LGN ROIs were calculated using Pearson’s partial correlations between each seed time series and each voxel in 4mm full-width-half-maximum smoothed resting-state images. Connectivity for left and right MGN and LGN TL fROIs was estimated separately, and the two connectivity maps for each geniculi pair were averaged together to produce a single image of bilateral MGN connectivity and a single image of bilateral LGN connectivity. This was repeated for alternative MGN and LGN ROIs generated from WFU PickAtlas, FreeSurfer thalamic segmentation, and THOMAS segmentation. To compare seed connectivity maps derived from TL MGN and LGN fROIs with those derived from each of their three alternatives, six seed connectivity difference images were then calculated for each participant: 1) TL MGN minus WFU MGN, 2) TL MGN minus FreeSurfer MGN, 3) TL MGN minus THOMAS MGN, 4) TL LGN minus WFU LGN, 5) TL LGN minus FreeSurfer LGN, and 6) TL LGN minus THOMAS LGN.

All Pearson’s partial correlations were calculated using the following nuisance parameters: band-pass filtered MPs (using the same 0.009-0.08 Hz band-pass filter that was applied to the RS time series), the squares of the band-pass filtered MPs, the derivatives of the band-pass filtered MPs, the squares of the derivatives of the band-pass filtered MPs, the white matter signal and its derivative, and the CSF signal and its derivative.

#### 2.9.4 Multi-Site Harmonization (ComBat)

ROI pair correlations and seed connectivity images were harmonized across SBU and NYSPI for site and scanner effects using ComBat (Fortin et al., 2018; Fortin et al., 2017; Johnson et al., 2007; Yu et al., 2018) (https://github.com/Jfortin1/ComBatHarmonization/) in MATLAB, using non-parametric adjustments and including demographic covariates. Demographic covariates were included in order to preserve biological variability in the sample while removing the variability associated with sites/scanners. Included covariates included age, gender, handedness, race, ethnicity, and parental socioeconomic status. PANSS Positive Scale, Negative Scale, and General Scale assessment scores were also included, to retain potential variability in auditory and visual perception, as well as other aspects of cognition and affect that might be associated with these measures.

Three participants were missing data for parental socioeconomic status and four participants were missing data for Positive and Negative Syndrome Scale assessments. Missing data were estimated in the full sample of participants who completed the study using SPSS Statistics (Version 29.0.0.0, Build 241, International Business Machines Corporation, Armonk, NY) as the aggregate median of 1,000 imputations, each with a maximum of 100 Markov chain Monte Carlo iterations, using linear regression models for scale variables and two-way interactions among categorical predictors. Singularity tolerance was set to 10^-12^; maximum case draws and maximum parameter draws were both set to 1,000. Missing data estimates determined via multiple imputation were only used as covariates in the ComBat harmonization procedure, and not in any other analyses.

### 2.10 Statistical Analysis

#### 2.10.1 Thalamic Localizer fROIs

As an exploratory analysis, significance was assessed for group-wide interhemispheric differences in TL-derived LGN and MGN fROI sizes using two-tailed Mann–Whitney–Wilcoxon rank-sum tests (Gibbons & Chakraborti, 2021; Hollander et al., 2014). Relationships between fROI size and connectivity with sensory cortex were then assessed, in order to determine whether differences in intrinsic functional connectivity that are observable in task coactivation maps could be systematically driving differences in the sizes of obtained LGN and MGN fROIs, or, conversely, whether differences in fROI sizes could be significantly driving estimates of measured connectivity with sensory cortex. Relationships between fROI sizes and MGN-AC connectivity were assessed using Kendall rank correlation coefficients (tau-b, two-tailed; Kendall, 1990; van Doorn et al., 2021).

#### 2.10.2. ROI Pair Correlation RSFC Analyses

The selectivity of time series extracted from TL MGN and LGN fROIs was evaluated using MGN-AC, MGN-VC, LGN-AC, and LGN-VC connectivity estimates. Using one-tailed paired t-tests, the following relationships were assessed, corrected for false discovery rate (Benjamini & Hochberg, 1995): 1) MGN selectivity for AC over VC (MGN-AC > MGN-VC), 2) AC selectivity for MGN over LGN (MGN-AC > LGN-AC), 3) LGN selectivity for VC over AC (LGN-VC > LGN-AC), and VC selectivity for LGN over MGN (LGN-VC > MGN-VC). One-tailed tests were used because the conclusion drawn from a non-significant result would not differ materially from the conclusion drawn from a significant two-tailed result in the unexpected direction; in both cases we would conclude that the TL task has failed to properly localize the appropriate thalamic nucleus.

Connectivity between TL fROIs and their appropriate sensory cortex (i.e., MGN-AC connectivity and LGN-VC connectivity) was then benchmarked against connectivity estimates calculated using alternative ROIs from WFU PickAtlas, the FreeSurfer thalamic nuclei segmentation, and the THOMAS segmentation, with the significance of differences assessed using two-tailed, paired t-tests.

#### 2.10.3 Seed Connectivity RSFC Analyses

Whole-brain TL MGN and LGN seed connectivity was assessed for significance using Permutation Analysis of Linear Models (PALM; Alberton et al., 2020; Winkler, Ridgway, et al., 2016; Winkler et al., 2014; Winkler, Webster, et al., 2016; Winkler et al., 2015), with a design matrix consisting of a single column of all 1s, using 10,000 sign-flips, built-in family-wise error rate correction (Holmes et al., 1996), and threshold-free cluster enhancement (TFCE; Smith & Nichols, 2009). All hypothesis testing was two-tailed (bi-sided), performed using two one-directional contrasts followed by multiple testing correction over contrasts (Alberton et al., 2020). This was repeated without TFCE as well, in order to improve the specificity of seed correlation results (at the expense of power). Exploratory analyses in PALM analyses were then conducted identically for difference images comparing seed connectivity between TL MGN and LGN fROIs, and those MGN and LGN ROIs obtained from WFU PickAtlas, FreeSurfer, and THOMAS. T-statistic maps obtained from PALM were thresholded at α *=* 0.05 after correction for family-wise error rate and over contrasts.

## 3. Results

### 3.1 Sensory Thalamic Localizer Auditory and Visual Thalamic fROIs (MGN and LGN)

A total of 52 participants (NYSPI n = 30; SBU n = 22) completed at least 2 runs each of the sensory TL and RS data. After evaluation of TL MGN and LGN fROIs, data from 47 participants (26 NYSPI; 21 SBU) were retained for subsequent analysis; fROIs that failed the QC procedure are shown in **Figure S4.** Sample demographics and clinical assessment data before and after exclusions for TL fROI failures are shown in **Table 1**.

**Table 1.**
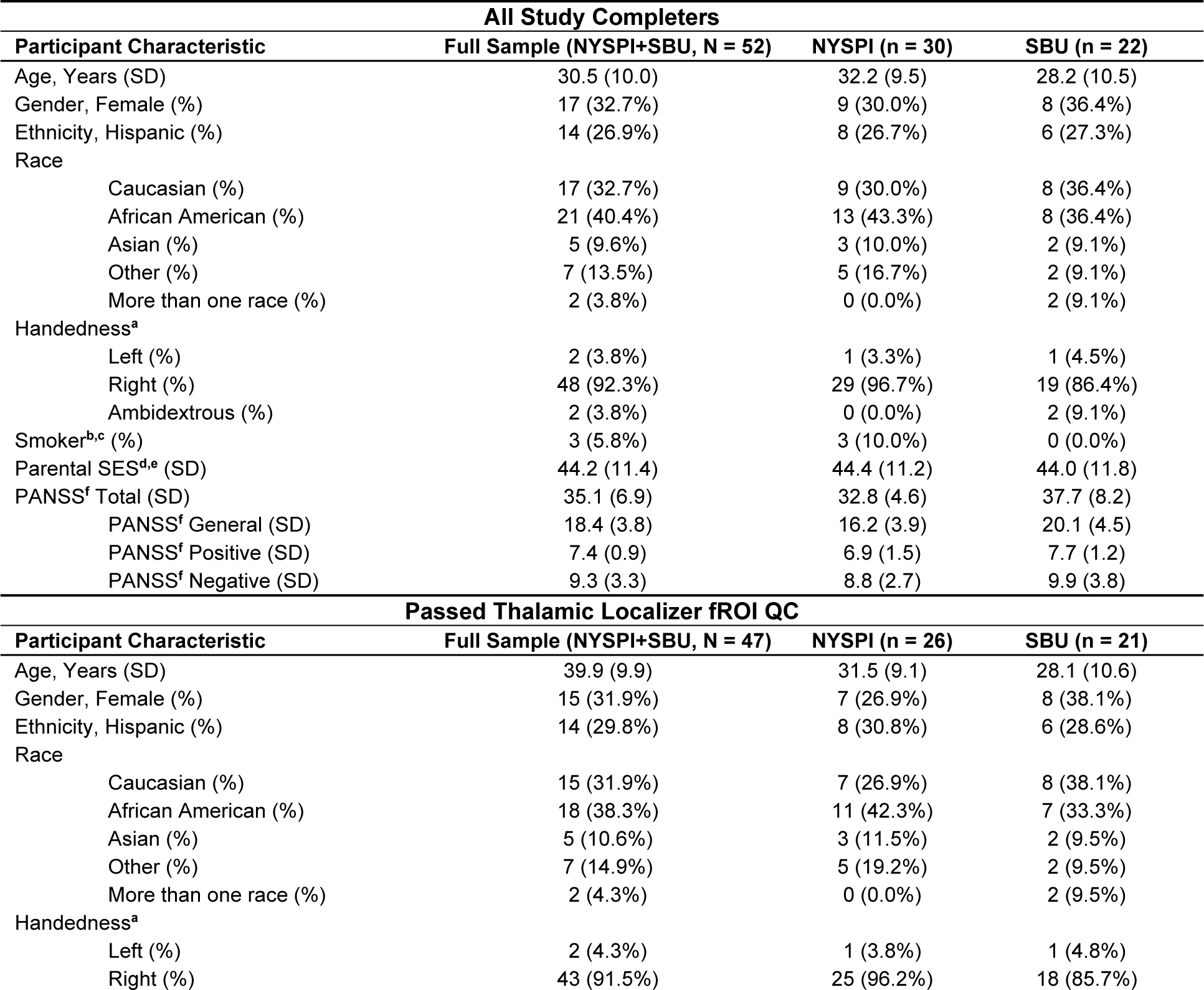
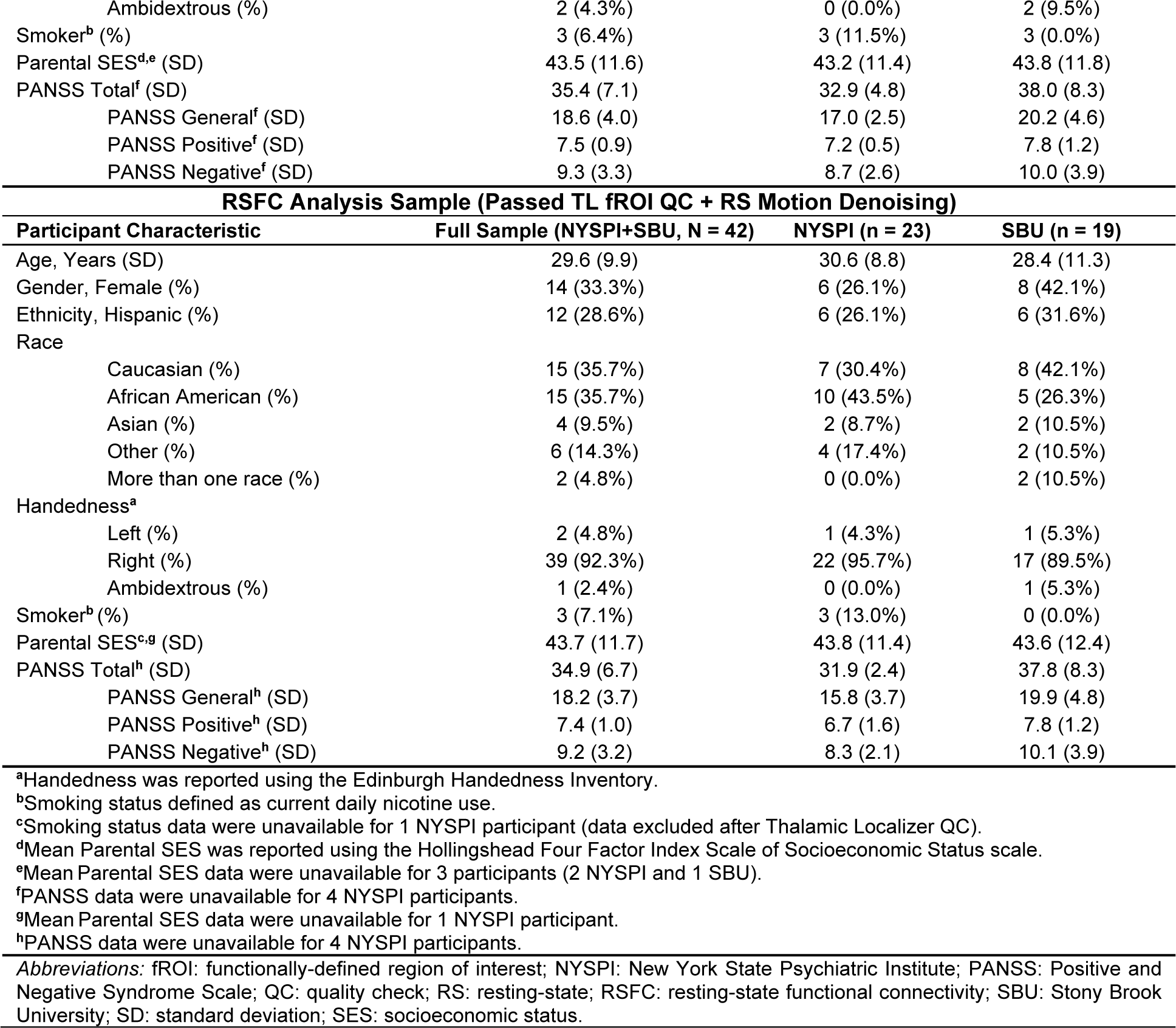
Demographic and clinical characteristics of study participants.

Group-wide probability density maps and corresponding violin plots of TL fROI sizes across participants are shown in **Figure 2** for left MGN **(Figure 2A-B)**, right MGN **(Figure 2C-D)**, left LGN **(Figure 2E-F)**, and right LGN **(Figure 2G-H).** Median left MGN and right MGN sizes were both 42 voxels, median left LGN size was 46 voxels, and median right LGN size was 50 voxels. LGN size was significant greater in the right than left hemisphere (p = 5.68 × 10^-4^), consistent with a recent finding using 7 Tesla structural quantitative MR imaging (Muller-Axt et al., 2021), but the interhemispheric difference in MGN size was not (p = 0.256). For visualization, probability density maps were thresholded (binarized) at 50% of the dataset, displayed together in **Figure 3**.

**Figure 2.**
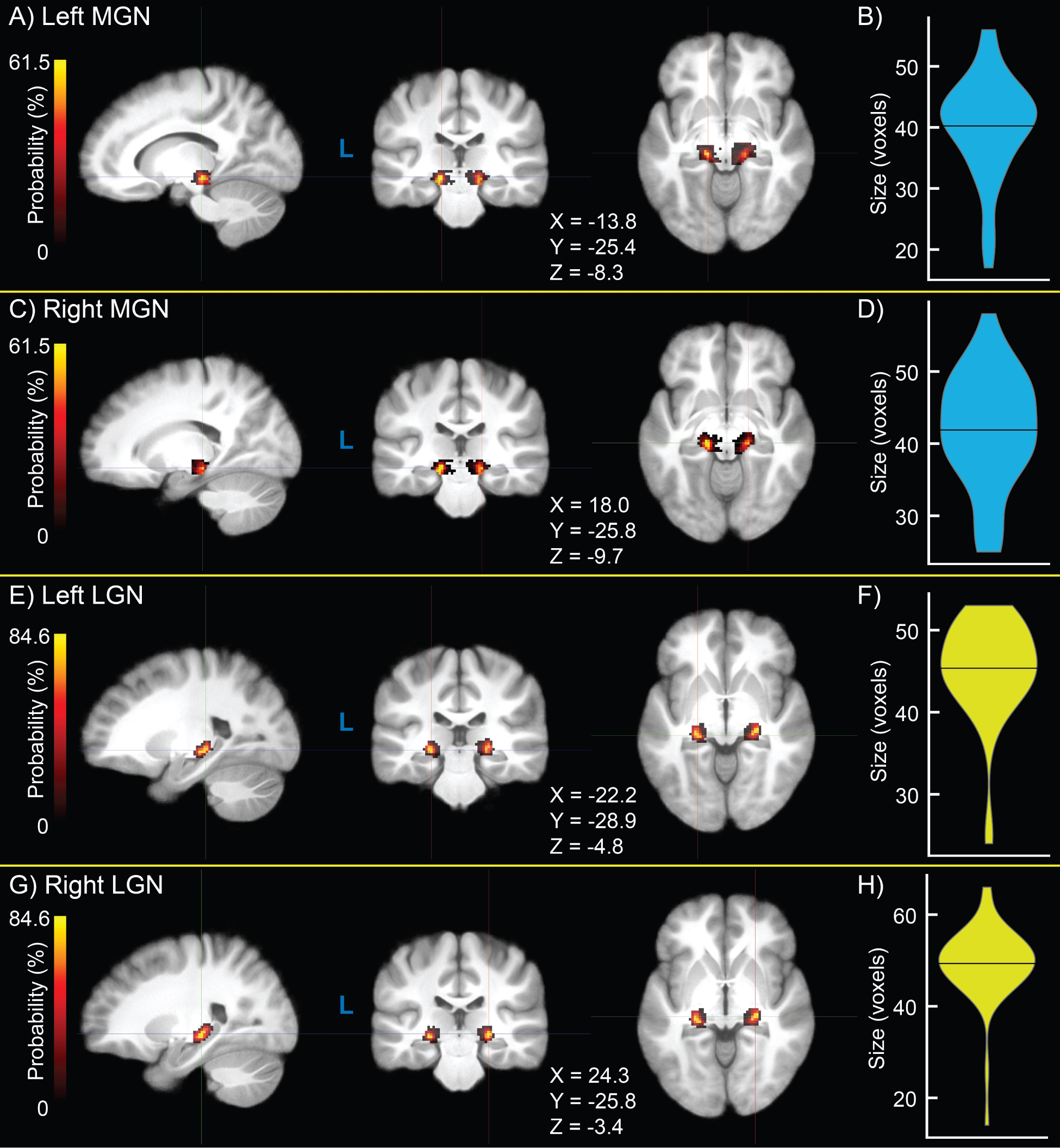
Group-level probability density maps of medial geniculate nucleus (MGN) and lateral geniculate nucleus (LGN) functionally-defined regions of interest (fROIs) obtained from the sensory thalamic localizer (TL) task **(A,C,E,G)** and violin plots of ROI size **(B,D,F,H)** for left MGN **(A,B)**, right MGN **(C,D)**, left LGN **(E,F)**, and right LGN **(G,H)**. Crosshair coordinates for each view are displayed with each set of images.

**Figure 3.**
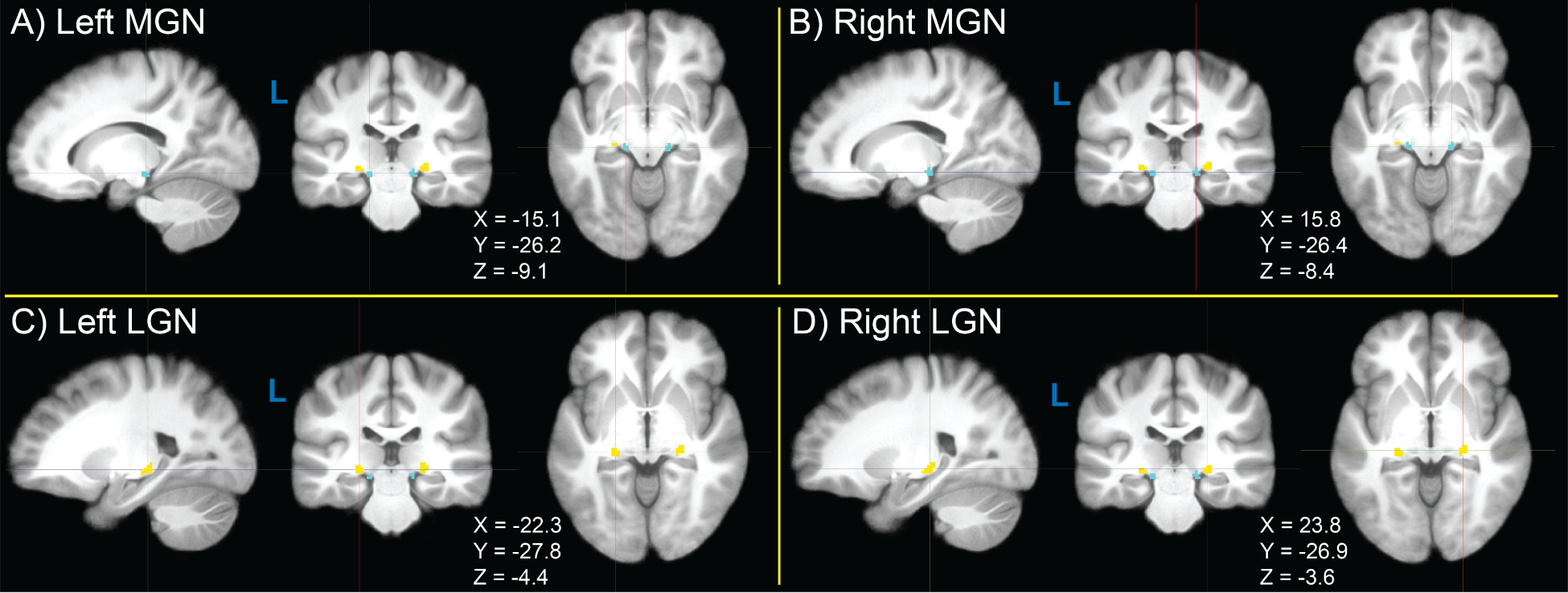
Thresholded (binarized) group-level probability density maps of left and right medial geniculate nucleus (MGN, blue) and lateral geniculate nucleus (LGN, yellow) functionally-defined regions of interest (fROIs) obtained from the sensory thalamic localizer (TL) task. Probability density maps for MGN and LGN were binarized by thresholding at 50% of participants. Image views and crosshairs are centered on the centroid of each fROI in Montreal Neurological Institute (MNI) 152 non-linear 6th-generation (MNI152NLin6), with crosshair coordinates shown for each set of images.

### 3.2 Resting-State Functional Connectivity Validation of Thalamic Localizer fROIs

Five participants (2 SBU, 3 NYSPI) were excluded from resting-state functional connectivity (RSFC) analyses due to insufficient data remaining after volume censoring, due to excessive participant motion. Descriptive statistics for the final RSFC analysis sample are shown in **Table 1**. An exploratory analysis assessing the association between the size of each hemisphere’s MGN and LGN size in each participant and connectivity with sensory cortex (AC and VC, respectively) using Kendall’s rank correlation coefficient are shown in **Table S1** (all p > 0.05).

Connectivity measured between MGN and LGN fROIs obtained from TL task analysis and AC and VC ROIs obtained from the FreeSurfer atlas (*Atlas_wmparc.2.nii.gz*) is shown in **Figure 4A**. Specificity measurements for MGN and LGN fROIs and their significance are shown in **Figure 4B** (all p < 0.05). Dice-Sørensen similarity coefficients between FreeSurfer-derived primary AC and primary VC ROIs and TL-derived AC and VC fROIs used as connectivity seeds for identifying MGN and LGN fROIs were 0.0533 and 0.115, respectively, indicating minimal bias in these analyses.

**Figure 4.**
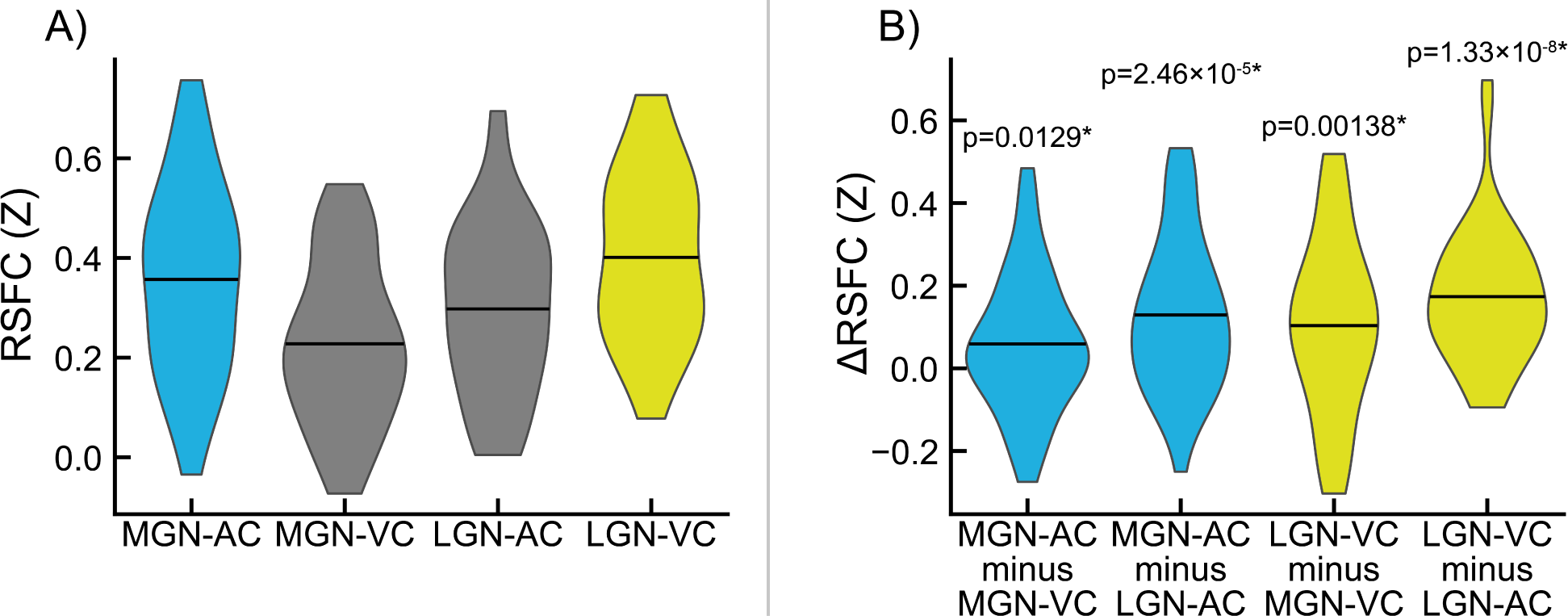
Evaluation of medial geniculate nucleus (MGN) and lateral geniculate nucleus (LGN) functionally-defined regions of interest (fROIs) obtained from the sensory thalamic localizer (TL) task. Pairwise resting-state function connectivity (RSFC) was evaluated between TL-derived geniculi and standard auditory cortex (AC) and visual cortex (VC) ROIs derived from the FreeSurfer atlas (*f*). **A)** Connectivity between MGN and AC (MGN-AC), MGN and VC (MGN-VC), LGN and AC (LGN-AC), and LGN and VC (LGN-VC). **B)** Selectivity of MGN and LGN ROIs for AC and VC, respectively, measured as within-participant differences in the RSFC values shown in panel A. MGN-AC minus MGN-VC quantifies MGN selectivity for AC over VC; MGN-AC minus LGN-AC quantifies AC selectivity for MGN over LGN. Likewise, LGN-VC minus MGN-VC quantifies VC selectivity for MGN over LGN, and LGN-VC minus LGN-AC quantifies LGN selectivity for VC over AC. P-values for significance of sample-wide mean difference are shown; asterisks (*) denote significance after false discovery rate correction (α = 0.05, one-tailed, one-sample t-test).

Whole-brain seed connectivity T-statistic maps are shown for TL MGN and LGN seed correlations, two-sided, thresholded at α *=* 0.05, and family-wise error rate corrected in **Figure S5.** Broadly distributed whole-brain connectivity is apparent in analyses using TFCE **(Figure S5A-B)**, with increased specificity apparent in voxel-wise (no TFCE) analyses **(Figure S5C-D).** In both images, significant connectivity is visible between MGN and AC, and between LGN and VC.

### 3.3 Resting-State Functional Connectivity Benchmarking of Thalamic Localizer fROIs

MGN-AC connectivity from MGN fROIs derived from the TL task are shown alongside MGN-AC connectivity from MGN ROIs derived from the WFU atlas (Maldjian et al., 2004; Maldjian et al., 2003), the FreeSurfer segmentation (Iglesias et al., 2018), and THOMAS segmentation (Su et al., 2019) in **Figure 5A**, and differences between MGN-AC connectivity estimates from TL fROIs and each of the other ROIs are shown in **Figure 5C**, along with P values for these connectivity differences. Similarly, LGN-VC connectivity estimates derived from TL task LGN fROIs are shown in **Figure 5B**, alongside LGN-VC connectivity estimates derived using WFU, THOMAS, and LGN ROIs. Differences in LGN-VC connectivity observed between TL LGN fROIs and the three alternatives are shown in **Figure 5D**. MGN-AC connectivity was significantly greater when using the TL task ROIs relative to the three alternatives (vs. WFU, p = 2.84×10^-9^; vs. THOMAS, p = 2.17×10^-4^; vs. FreeSurfer, p = 3.36×10^-2^). TL LGN-VC connectivity was observed to be greater relative to the three alternatives, significantly versus WFU (p = 9.00×10^-14^) and THOMAS (p = 6.10×10^-8^), but not significantly versus FreeSurfer (p = 0.584).

**Figure 5.**
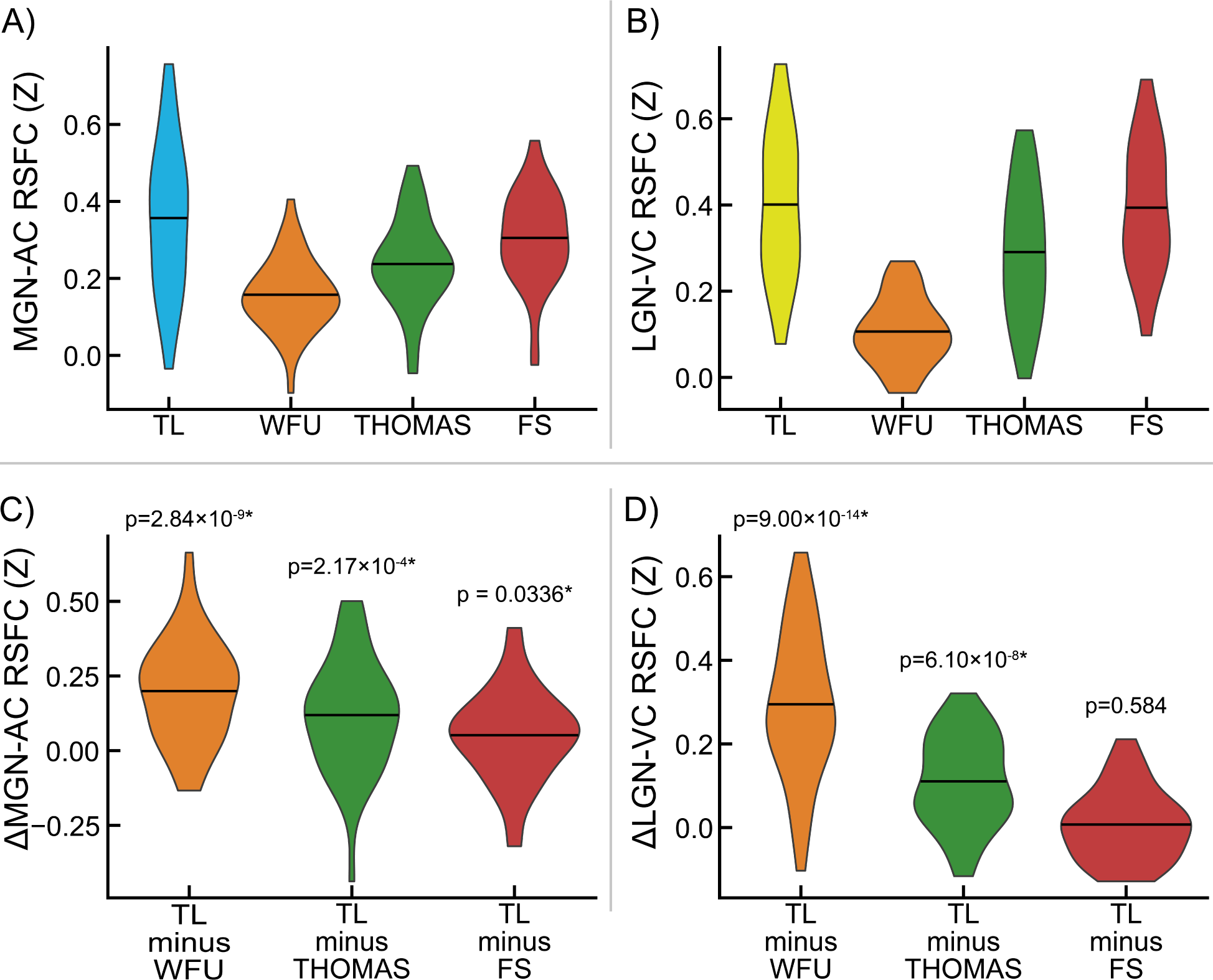
Benchmarking of medial geniculate nucleus (MGN) and lateral geniculate nucleus (LGN) functionally-defined regions of interest (fROIs) obtained from the sensory thalamic localizer (TL) task through resting-state functional connectivity (RSFC) with auditory cortex (AC) and visual cortex (VC) ROIs, respectively. AC and VC regions of interest (ROIs) were obtained from the FreeSurfer (FS) Desikan-Killiany atlas (*Atlas_wmparc.2.nii.gz*). MGN and LGN derived from the TL task are evaluated relative to those derived from 1) MNI space atlas ROIs from the Wake Forest University (WFU) PickAtlas, 2) THOMAS segmentation applied to each subject’s T1w image, and 3) the FreeSurfer thalamic segmentation applied to each subject’s T1w. Panel **A** shows MGN-AC RSFC using the TL MGN ROI alongside the three named alternatives; panel **B** shows the within-participant differences between MGN-AC connectivity from the TL-derived MGNs and the three alternatives (i.e., differences between TL and other values in panel A). Likewise, panel **C** shows LGN-VC RSFC using the TL LGN ROI alongside alternatives derived from WFU, THOMAS, and FS; panel **D** shows the within-participant differences between TL connectivity and the three alternatives shown in panel **C**. P-values for significance of sample-wide mean difference are shown; asterisks (*) denote significance (α < 0.05, two-tailed, one-sample t-test).

Exploratory analyses assessing for differences in seed connectivity between TL MGN and LGN fROIs, and the three evaluated alternatives (WFU, THOMAS, and FreeSurfer), are shown in **Figure 6**. Significant whole-brain increases in connectivity brain are visible relative to WFU ROIs **(Figure 6, panels A and D).** TL MGN fROIs show increased connectivity with areas of primary AC and superior temporal gyrus relative to all alternative ROIs **(Figure 6A-C).** Similarly, TL LGN fROIs show increased connectivity compared to the WFU and THOMAS ROIs across the brain, with particularly increased connectivity with regions of primary VC around the calcarine sulcus **(Figure 6D-E).** No significant differences in TL LGN seed connectivity and FreeSurfer seed connectivity were found after correction for family-wise error rate.

**Figure 6.**
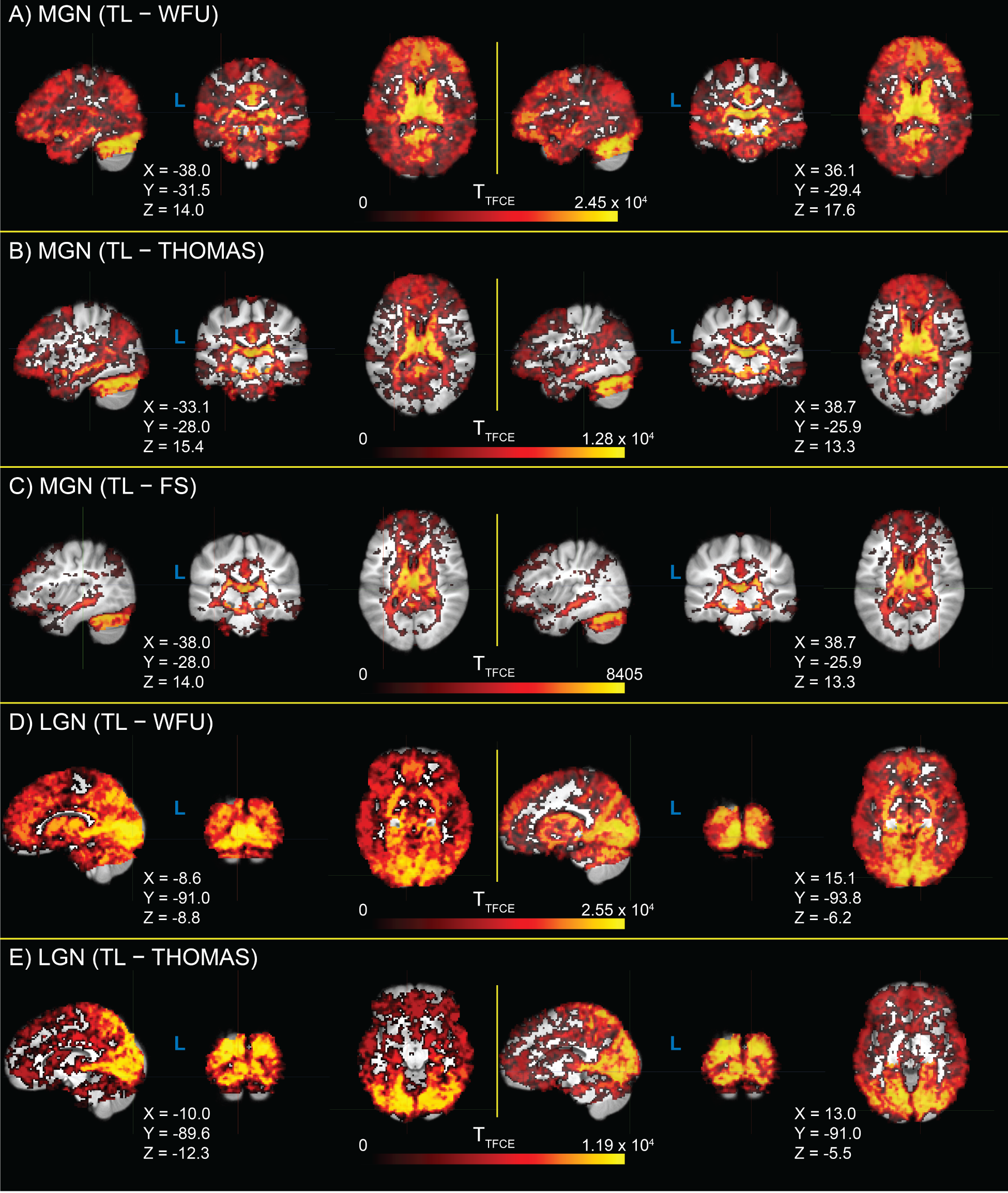
Whole-brain T-statistic maps for differences in seed connectivity between thalamic localizer (TL) derived medial geniculate nucleus (MGN) and lateral geniculate nucleus (LGN) functionally-defined regions of interest (fROIs), relative to seed connectivity calculated from three alternative regions of interest (ROIs): Wake Forest University PickAtlas (WFU), the Thalamus Optimized Multi Atlas Segmentation (THOMAS), and the FreeSurfer thalamic nuclei segmentation (FS). T-statistics are thresholded at α = 0.05 (two-tailed, family-wise error rate corrected). Image views are centered on either auditory cortex (panels **A-C**) or visual cortex (panels **D** and **E**) of each hemisphere, with crosshair coordinates for each view shown. A panel for LGN TL − FS T-statistics is not shown, as there were no significant voxels.

## 4. Discussion

Here, we detail a sensory thalamic localizer (TL) fMRI task (design shown in **Figure 1**) capable of localizing auditory and visual thalamic nuclei in individual human participants (shown in **Figure 2** and **Figure 3)**. Building off of past work (Chen et al., 1998; Denison et al., 2014; Jiang et al., 2013; Kastner et al., 2004), in which localizer tasks were utilized in order to identify either medial or lateral geniculate nuclei, here we show identification of both MGN and LGN through the use of a single task. Additionally, this task provides estimates of these thalamic nuclei in 16 minutes 12 seconds total, across 4 runs (of 3 minutes 46 seconds each), which is substantially less time than in prior work. For example, a single set of auditory thalamic fROIs was acquired in 32 minutes, across 6 runs (of 5 minutes and 20 seconds each) (Jiang et al., 2013). This is likely at least partially due to improvements in data acquisition: data here were acquired using 3 Tesla MR scanners, using a head coil with 32 or more channels (i.e., 32-channel at NYSPI and 48 head coil channels in the 64-channel head-and-neck coil at SBU), a high-resolution simultaneous-multi-slice (multiband) acquisition sequence (Moeller et al., 2010), and clustered-sparse temporal acquisition (Schmidt et al., 2008; Yang et al., 2000; Zaehle et al., 2007) of 3 volumes per cluster.

Additionally, improvements in the ability to resolve the location of auditory and visual thalamic fROIs is potentially enabled in part due to novel elements of the task design, in which a contrast between auditory and visual stimuli was used to robustly detect responsive regions around auditory and visual cortices in each subject, and use these task time series to find thalamic voxels (within specified search regions) that are responsive to either, but not to both. The contrast produced by alternating task conditions serves as a sensory control condition, in which circuits involved in the conscious perception of stimulation generally are engaged in both conditions. Furthermore, this design increases the specificity of obtained thalamic nuclei: auditory and visual thalamic fROIs are optimized to represent regions specific to their sensory modality, and not general features of perception or cognition. This may be critical in studies that aim to relate human perceptual abnormalities to dysfunction in circuits specifically associated with either auditory or visual processing. We validated this specificity using resting-state functional connectivity (RSFC) data **(Figure 4)** collected from the same participants.

Finally, we compared fROIs obtained from the TL task to three alternative sources of thalamic ROIs **(Figure 5)**: an atlas-based ROI from the Wake Forest University PickAtlas Toolbox (Maldjian et al., 2004; Maldjian et al., 2003), the Thalamus Optimized Multi Atlas Segmentation (THOMAS; Su et al., 2019) and the FreeSurfer thalamic nuclei segmentation (Iglesias et al., 2018). We found significantly greater connectivity with auditory cortex from TL MGN fROIs relative to the evaluated alternatives, and significantly greater connectivity with visual cortex from TL LGN fROIs relative to the WFU and THOMAS ROIs; average connectivity from the FreeSurfer LGN ROI to visual cortex was slightly increased, but not significantly so. This may be due in part to the fact that FreeSurfer MGN ROIs are utilized during TL task analysis as the starting point for building search regions for these fROIs, and may additionally represent that priors for LGN locations in the FreeSurfer segmentation are typically well-placed for detecting BOLD activation from visual thalamus. This pattern is also visible in exploratory whole-brain seed correlation analyses exploring regions showing significant differences in connectivity from TL MGN and LGN fROIs relative to the three aforementioned alternative ROIs **(Figure 6).**

### 4.1 Caveats and Limitations

One critical aspect of fROIs derived from the TL task described here is that these fROIs purport to represent the profile of the hemodynamic signature of geniculate nuclei’s neural responses during task stimulation as detected in BOLD echo planar images, including the point spread functions of the hemodynamic response and T2* contrast (Parkes et al., 2005; Shmuel et al., 2007; Ugurbil, 2016; Ugurbil et al., 2013; Vu et al., 2017), after pre-processing and post-processing (including spatial smoothing). Thus they may not correspond fully to anatomically expected sizes observed in human tissue post-mortem, or *in vivo* using anatomical (as opposed to functional) imaging techniques (Garcia-Gomar et al., 2019; Muller-Axt et al., 2021; Oishi et al., 2023; Sitek et al., 2019; Tourdias et al., 2014). When using these fROIs in BOLD fMRI data analysis, this could be a substantial advantage, since they may better map onto specific spatial features of the thalamic hemodynamic response and its profile in BOLD images, which can be impacted by imperfectly corrected susceptibility distortions, errors in spatial normalization to MNI template space, and other errors in localization. However, depending on the application, this advantage may be a flaw, and other techniques may prove superior; for example, when attempting to evaluate specific features of individual anatomy that are too fine to be reliably detected with standard whole-brain 3 Tesla BOLD imaging techniques. Likewise, aspects of human perceptual processing that occur at faster timescales than the Nyquist frequency enabled by BOLD fMRI imaging techniques (e.g., 800 ms) are not detectable using fMRI, except through summation into a slower, aggregate hemodynamic response.

Additionally, TL task-derived fROIs aim to achieve a degree of specificity to sensory modality, which could preclude their use for aspects of perception that involve multisensory integration or context-dependent modulation, including the medial and dorsal subnuclei of the MGN (Lee, 2015; Wepsic, 1966). While this does not appear to be a significant issue given the smoothness of 3T BOLD fMRI, especially after Gaussian smoothing, and the relatively large voxel size of 2 mm (isotropic), this could require modification if adapting this task for more precise mapping of geniculi using higher field strengths or more specialized sequences.

The TL task requires dedicated data acquisition of slightly more than 15 minutes total; this is in addition to standard T1w and T2w sequences, which can be used to obtain ROIs from anatomical segmentations. Future work systematically exploring the impact of scan time on fROI quality is warranted in order to aid in balancing needs for data quality against the feasibility of adding the task to a scan protocol. In this study, task data acquired from 5 of 52 (9.6%) participants did not produce a full set of four valid thalamic fROIs, which may be a concern in studies with limited sample sizes or variable data quality, and the extent to which additional data collection could have reduced this failure rate is unknown. As this study did not include the evaluation of test-retest reliability of fROIs obtained from the sensory TL task, future work should aim to quantify this by obtaining multiple sessions of sensory TL task data for each participant and explore the relationship between reliability measures and scan time.

Finally, the task code is written in Presentation, and the analysis code in MATLAB. These are proprietary software tools that require paid licenses to use. Future work exploring the adaptation of the task code to utilize free alternatives, such as Python (Python Software Foundation, Wilmington, DE) and PsychoPy (Peirce et al., 2019; Peirce, 2007, 2008), would allow for greater accessibility to researchers. As this software is open-source and released under the GNU General Public License version 3, it may be translated to other platforms and programming languages, as well as extended or adapted for other purposes as desired by the wider research community.

### 4.2 Conclusion

This work details the development, implementation, validation, and benchmarking of a sensory thalamic localizer (TL) BOLD fMRI task capable of detecting auditory and visual thalamic nuclei in individual participants. Using resting-state functional connectivity analyses, we show that auditory and visual thalamic functionally-defined regions of interest (fROIs) obtained from the sensory TL task are modality specific, and benchmark them against existing atlas-and T1w segmentation-based alternatives. This task produces estimated thalamic medial and lateral geniculate fROIs suitable for resting-state functional connectivity (RSFC) studies and may be added to fMRI acquisition protocols with less than 16 minutes of dedicated additional acquisition time. The TL task, implemented in Presentation, and analysis code, implemented in MATLAB, are both publicly available to the research community.

## 5 Data and Code Availability

The sensory Thalamic Localizer Presentation task code (Williams et al., 2024) is available from GitHub at https://github.com/CNaP-Lab/Sensory-Thalamic-Localizer <u>and</u> the Neurobehavioral Systems Archives of Neurobehavioral Experiments and Stimuli at http://www.neurobs.com/ex_files/expt_view?id=302. The MATLAB task analysis code for producing MGN and LGN fROIs from acquired BOLD fMRI task data and Presentation task logs is additionally available from GitHub at https://github.com/CNaP-Lab/Sensory-Thalamic-Localizer. All software associated with this manuscript is released under the GNU General Public License version 3. Note that the sensory Thalamic Localizer task is designed to be deployed on a system with a display refresh rate of 60 Hz. Data acquired from human participants used in the analyses detailed in this manuscript are available upon request from the corresponding author through a formal data sharing agreement.

## 6 Author Contributions

**Conceptualization:** G.H., A.A-D., and J.X.V.S.; **Data Curation:** J.C.W., P.N.T., Z.J.Z., E.B.S-F., D.T.P., N.K.H., S.K.A., G.H., and J.X.V.S.; **Formal Analysis:** J.C.W., P.N.T., and J.X.V.S.; **Funding Acquisition:** J.C.W., A.A-D., G.H., and J.X.V.S.; **Investigation:** J.C.W., P.N.T., Z.J.Z., E.B.S-F., D.T.P., N.K.H., S.K.A., A.A-D., G.H., and J.X.V.S.; **Methodology:** J.C.W., P.N.T., G.H., and J.X.V.S.; **Project Administration:** J.C.W., N.K.H., A.A-D., G.H., and J.X.V.S.; **Resources:** A.A-D., G.H., and J.X.V.S.; **Software:** J.C.W., P.N.T., S.K.A., and J.X.V.S.; **Supervision:** N.K.H., A.A-D., G.H., and J.X.V.S.; **Validation:** J.C.W., S.K.A, G.H., and J.X.V.S.; **Visualization:** J.C.W., P.N.T., D.T.P., and J.X.V.S.; **Writing – Original Draft:** J.C.W.; **Writing – Review & Editing:** J.C.W., P.N.T., Z.J.Z., A.A-D., G.H., and J.X.V.S.

## 7 Funding

Research reported in this publication was supported by the National Institute of Mental Health of the National Institutes of Health (NIH) under award numbers K01MH107763 to J.X.V.S., K23MH101637 to G.H., R01MH109635 to A.A-D., and F30MH122136 to J.C.W. J.C.W. was also supported by an NIH Research Supplement to Promote Diversity in Health-Related Research (3R01MH120293-04S1) and by the Stony Brook University Medical Scientist Training Program (NIH Award No. T32GM008444; Principal Investigator: Dr. Michael A. Frohman). P.N.T. was supported by a Stony Brook University Department of Biomedical Engineering Graduate Assistance in Areas of National Need Fellowship (United States Department of Education Award No. P200A210006; Director: Dr. David Rubenstein) and the Stony Brook University Scholars in Biomedical Sciences Program (NIH Award No. T32GM148331; PI: Dr. Styliani-Anna [Stella] E. Tsirka). The Stony Brook high-performance SeaWulf computing system was supported by National Science Foundation (NSF) Award Nos. 1531492 (PI: Dr. Robert Harrison; co-PI: Dr. Yuefan Deng) and 2215987 (PI: Dr. Robert Harrison; co-PIs: Dr. Yuefan Deng, Dr. Eva Siegmann, and David Cyrille), and matching funds from the Empire State Development’s Division of Science, Technology and Innovation (NYSTAR) program contract C210148. Magnetic resonance imaging at Stony Brook University was performed at the Social, Cognitive, and Affective Neuroscience (SCAN) Center, supported by NSF Award No. 0722874 (PI: Dr. Turhan Canli). The content is solely the responsibility of the authors and does not necessarily represent the official views of the NIH, NSF, or NYSTAR.

## 8 Declaration of Competing Interests

Anissa Abi-Dargham received consulting fees and/or honoraria from Sunovion Pharmaceuticals Inc., Otsuka Pharmaceutical Co., Ltd., Merck & Co., Inc., Neurocrine Biosciences, Inc., F. Hoffmann-La Roche AG, and C.H. Boehringer Sohn AG & Co. KG. Anissa Abi-Dargham holds stock options in Herophilus, Inc. and in Terran Biosciences, Inc. All other authors declare that they have no known competing financial interests or personal relationships that could have influenced or appear to have influenced the work reported in this paper.

## 9 Acknowledgements

The authors would like to thank Jaeyop Jeong, Sam R. Luceno, Srineil Nizambad, and Yash Patel for contributions to fMRI data preprocessing, Kelly Bobchin for assistance with data organization, and Tram Ngoc Bao Nguyen for helpful discussions. Computing resources and technical assistance were provided by Stony Brook Medicine Research Computing, with substantial support from Allen Zawada and James Xikis. Access to and technical support for the high-performance SeaWulf computing system was provided by Stony Brook Research Computing and Cyberinfrastructure and the Institute for Advanced Computational Science at Stony Brook University, with notable support from Fırat Coşkun, Daniel Wood, and David Carlson.

## Notes

http://www.neurobs.com/ex_files/expt_view?id=302

https://github.com/CNaP-Lab/Sensory-Thalamic-Localizer

### Summary of Updates

Expanded discussion and minor additions to methods. Removed excess abbreviations for readability.

