## Supplementary Material for "Functional Localization of the Human Auditory and Visual Thalamus Using a Thalamic Localizer Functional Magnetic Resonance Imaging Task"

**Affiliations**

**Address:** 101 Nicolls Rd., Health Sciences Center T10-087J, Stony Brook, NY 11794

**Short/running title:** fMRI Localization of the Human Auditory and Visual Thalamus

### Supplementary Materials and Methods

#### *Inclusion and Exclusion Criteria*

At New York State Psychiatric Institute (NYSPI), inclusion criteria were: 1) ages 18-55, 2) absence of any current Diagnostic and Statistical Manual of Mental Disorders, Fourth Edition, Text Revision (DSM-IV-TR; American Psychiatric Association, 2000) Axis-I psychiatric disorder, and 3) fluent in spoken English. NYSPI exclusion criteria were: 1) current substance use disorders; 2) current major affective episode; 3) history of neurological disorders, including head trauma with loss of consciousness, epilepsy, or other clinically significant disorders of the central nervous system, intellectual disability, hearing impairment, or unstable severe medical conditions; 4) claustrophobia; 5) metal implants or paramagnetic objects contained within the body; and 6) current pregnancy.

At Stony Brook University (SBU), inclusion criteria were: 1) ages 18-55; 2) negative urine toxicology; 3) hearing threshold within normal limit (up to 25 decibels Hearing Level from 250 Hz through 2 kHz, bilaterally); 4) speech recognition for monosyllabic words equal to or better than 80% correct. Exclusion criteria at SBU were: 1) current or past, or family history of, psychiatric illness, including alcohol or substance abuse, except nicotine use disorder; 2) clinically significant medical or neurological illness; 3) pregnancy or lactation; 4) lack of effective birth control; 5) presence of metallic objects in the body; 6) current, past, or anticipated exposure to radiation in the workplaces, or participation in nuclear medicine procedures elsewhere; 7) current aural or neurological disease; 8) more than one risk factor for coronary disease; 9) insulin-dependent diabetes; 10) history of significant cardiovascular illness; 11) stage 2 hypertension (blood pressure greater than 140 mm Hg systolic or 90 mm Hg diastolic); 12) clinically significant brain abnormalities; 13) prisoners; 14) severe claustrophobia; and 15) any previous severe reaction to amphetamine. Psychiatric diagnoses at SBU were established according to the Diagnostic and Statistical Manual of Mental Disorders, Fifth Edition (DSM-5; American Psychiatric Association, 2013).

Positive pregnancy screens resulted in cancellation of the magnetic resonance (MR) scan session and exclusion of the participant from further participation in the study while pregnant. In the event of a positive urine toxicology test result, MR scanning was aborted and rescheduled at the discretion of participants and study personnel.

### Supplementary Tables

**Table S1.** Correlations (Kendall rank correlation coefficients) between the size of TL-derived MGN and LGN fROIs and connectivity with sensory cortex

| ROI | RSFC ROI Pair | Kendall's tau-c | P-Value (2-tailed) |
|---|---|---|---|
| Left MGN | Left MGN - Left AC | 0.063 | 0.572 |
| | Left MGN - Right AC | 0.092 | 0.409 |
| Right MGN | Right MGN - Right AC | 0.039 | 0.728 |
| | Right MGN - Left AC | 0.006 | 0.965 |
| Left LGN | Left LGN - Left VC | 0.104 | 0.350 |
| | Left LGN - Right VC | 0.113 | 0.307 |
| Right LGN | Right LGN - Right VC | 0.157 | 0.155 |
| | Right LGN - Left VC | 0.154 | 0.161 |

Abbreviations: AC: Auditory Cortex; fROI: functionally-defined region of interest; LGN: Lateral Geniculate Nucleus; MGN: Medial Geniculate Nucleus; TL: thalamic localizer; VC: Visual Cortex.

### Supplementary Figures

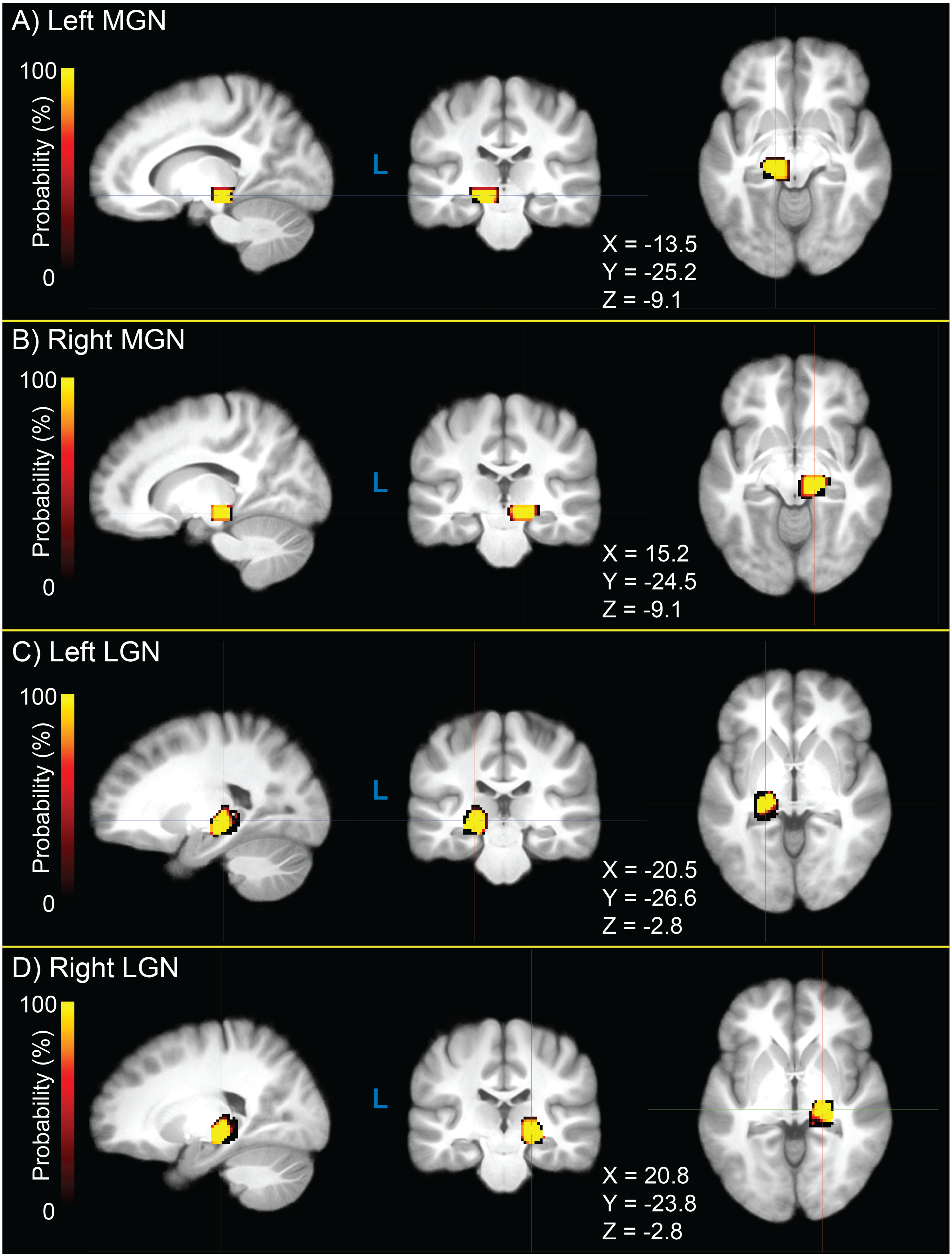

**Figure S1.** Thalamic search regions (TSRs) for medial geniculate nucleus (MGN, panels **A** and **B**) and lateral geniculate nucleus (LGN, panels **C** and **D**) used to identify functionally-defined regions of interest (fROIs) using data acquired during the sensory thalamic localizer (TL) functional magnetic resonance imaging (fMRI) task.

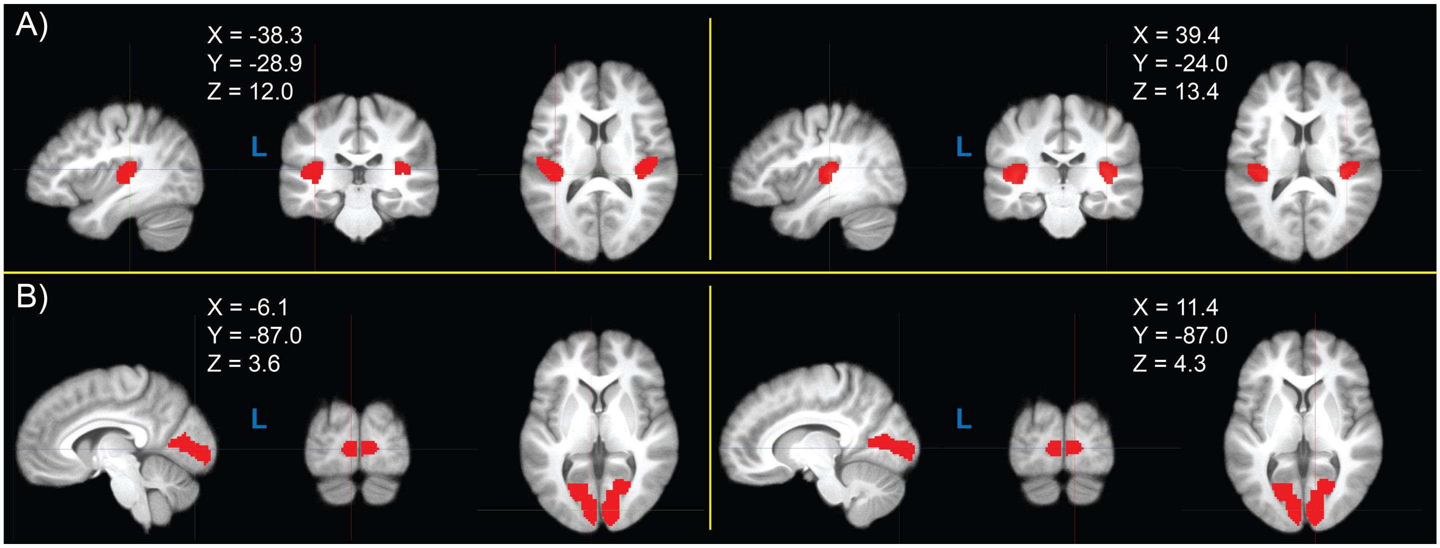

**Figure S2.** Regions of interest (ROIs) of auditory cortex (AC; transverse temporal gyrus; panel **A**) and visual cortex (VC; paracalcarine gray matter; panel **B**) used for resting-state connectivity, derived from the FreeSurfer Desikan-Killiany atlas parcellation (*Atlas_wmparc.2.nii.gz*). Crosshair coordinates for each view are displayed with each set of images.

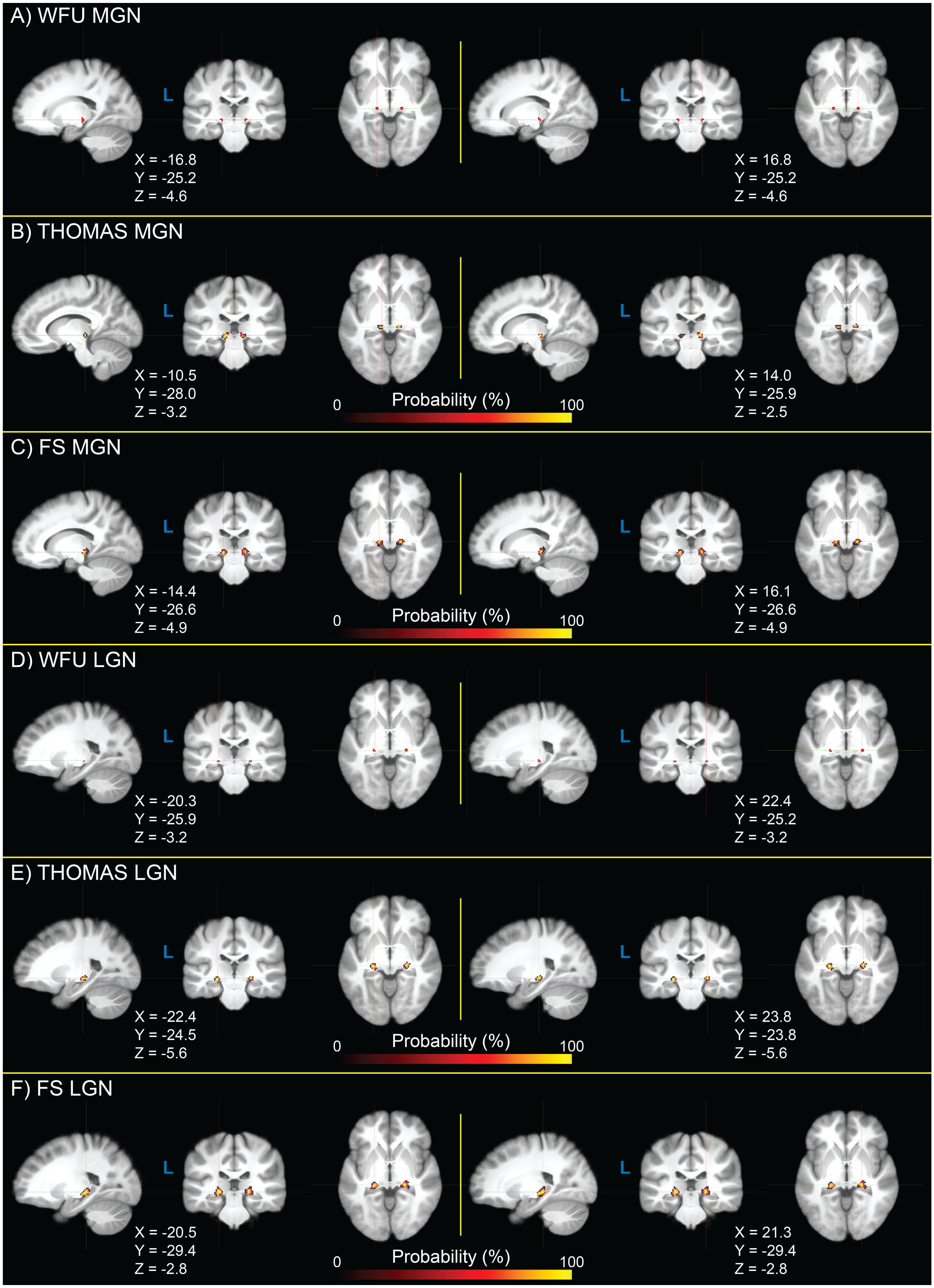

**Figure S3.** Medial geniculate nucleus (MGN) and lateral geniculate nucleus (LGN) regions of interest (ROIs) derived from Wake Forest University (WFU) PickAtlas **(A,D)**, the THOMAS T1w segmentation **(B,E)**, and the FreeSurfer (FS) thalamic segmentation **(C,F)**. WFU ROIs are identical across participants, as they are atlas-based. THOMAS and FS ROIs are derived from T1w image segmentations, which vary across participants, and are thus shown as probability density maps across the study sample. Crosshair coordinates for each view are displayed with each set of images.

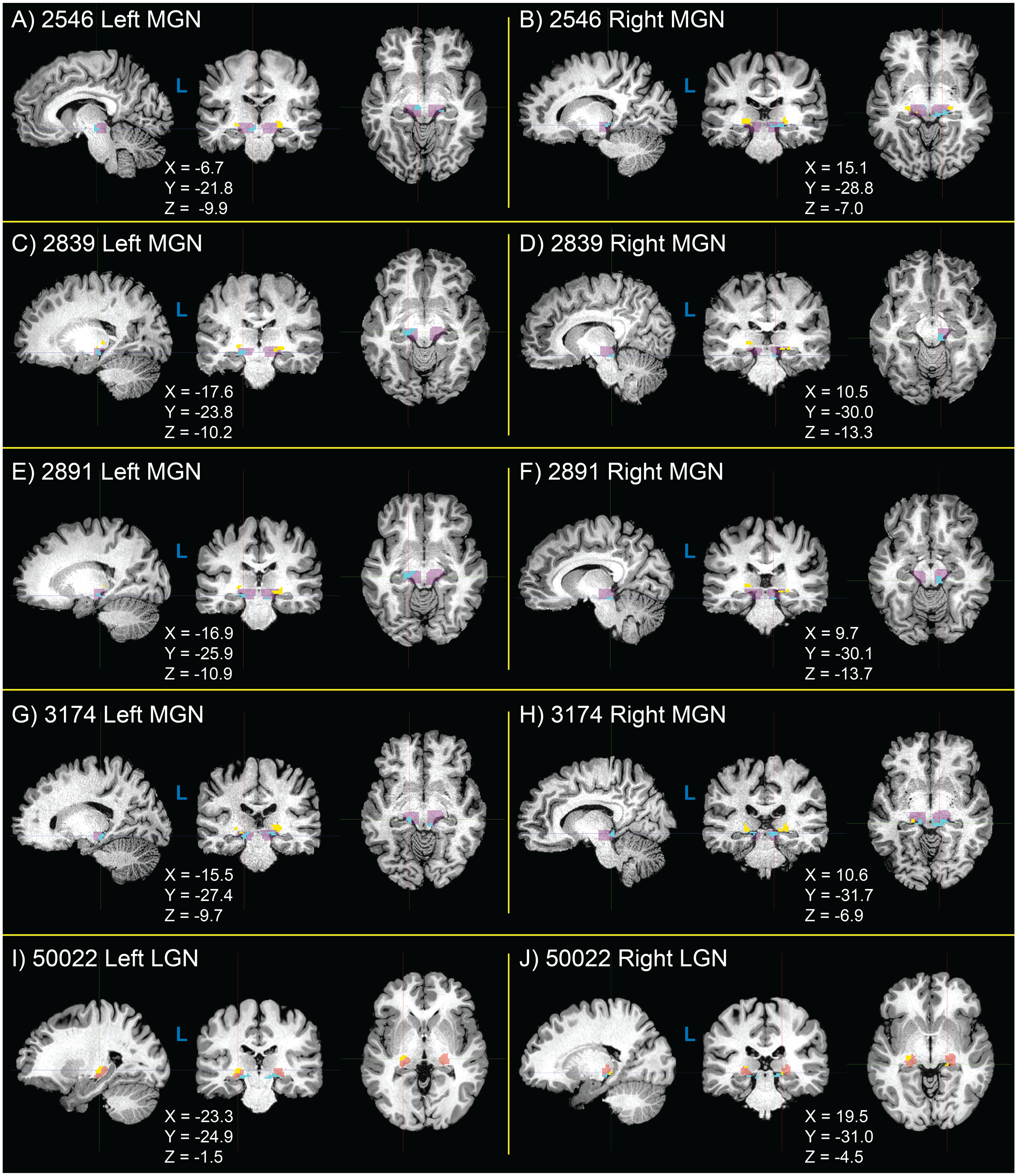

**Figure S4.** Functionally-defined regions of interest (fROIs) for medical geniculate nucleus (MGN, panels **A-H**) and lateral geniculate nucleus (LGN, panels **I-J**) obtained from the sensory thalamic localizer (TL) task that failed quality check (QC) procedures. MGN fROIs are shown in cyan; LGN fROIs are shown in yellow. MGN thalamic search regions (MGN-TSRs) are shown in purple, and LGN thalamic search regions (LGN-TSRs) are shown in red. In each case of an fROI QC failure, the fROI from both hemispheres are shown for comparison. The noted causes of fROI QC failures are as follows: **A,B)** excessive anterior-posterior (AP) MGN asymmetry; **C,D)** excessive MGN AP asymmetry and apparent signal from midbrain in right MGN, likely from inferior colliculus (IC); **E,F)** excessive MGN AP asymmetry and apparent midbrain (IC) signal in right MGN; **G,H)** MGN AP and left-right (LR) asymmetry, potentially due to midbrain (IC) signal in right MGN; **I,J)** excessive asymmetry between left and right LGN position and size, potentially due to capture of extra-thalamic signal.

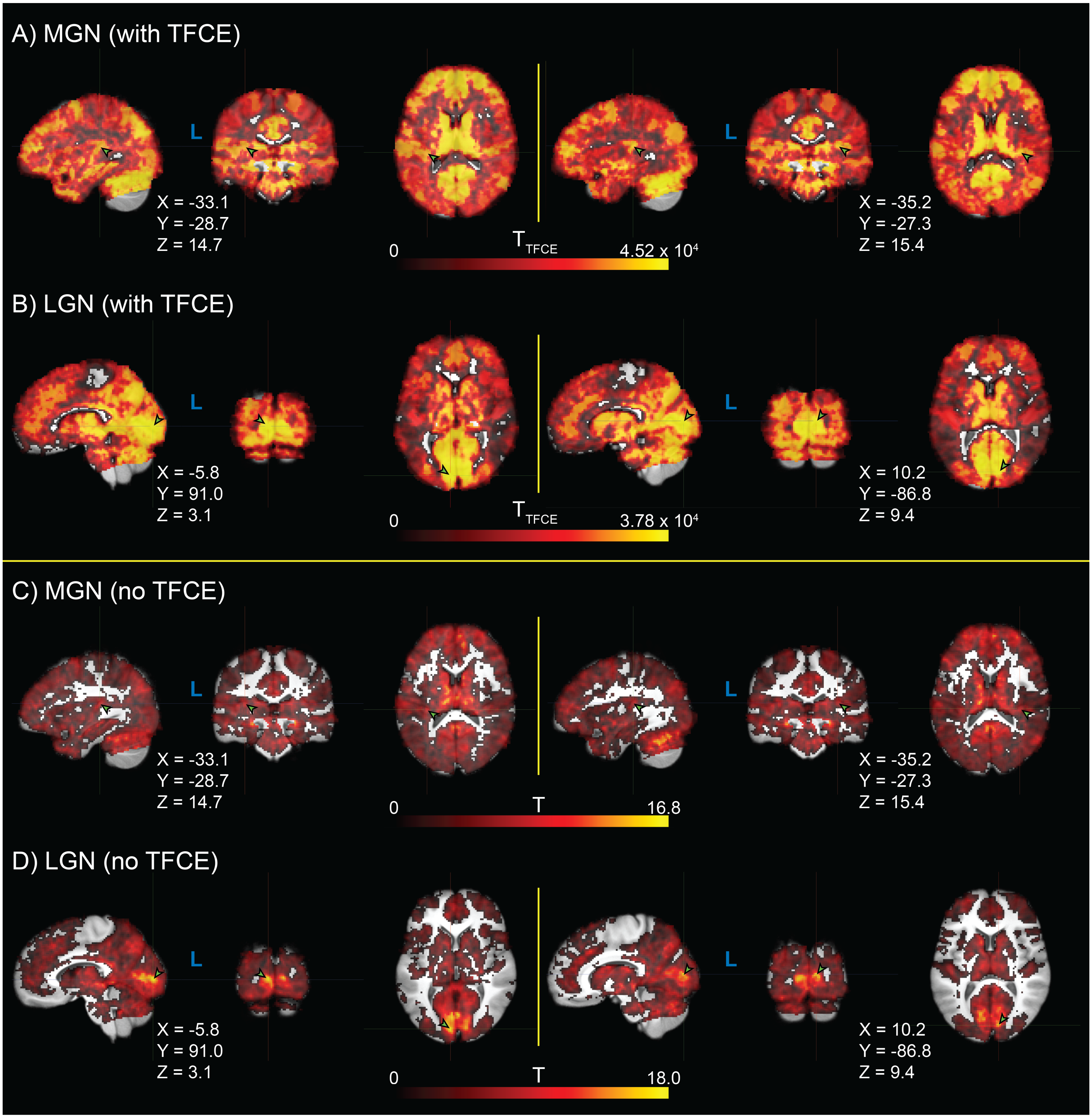

**Figure S5.** Whole-brain seed connectivity T-statistics from medial geniculate nucleus (MGN) **(A,C)** and lateral geniculate nucleus (LGN) **(B,C)** functionally-defined regions of interest (fROIs) derived from the sensory thalamic localizer (TL) task. Results are shown both with threshold-free cluster enhancement (TFCE) **(A,B)** and without TFCE **(C,D)**. T-statistics are thresholded at $\alpha = 0.05$ (two-tailed and family-wise error rate corrected). Image views are centered on either primary auditory cortex (transverse temporal gyrus; panels **A** and **C**) or visual cortex (paracalcarine gray matter; panels **B** and **D**) of each hemisphere highlighted by green arrows with black border. Crosshair coordinates for each view are shown for each set of images.
